# PDX3 is important for carbon/nitrogen balance in Arabidopsis associated with distinct environmental conditions

**DOI:** 10.1101/2022.12.06.519276

**Authors:** Priscille Steensma, Marion Eisenhut, Maite Colinas, Laise Rosado-Souza, Alisdair R. Fernie, Andreas P. M. Weber, Teresa B. Fitzpatrick

**Author notes:** Teresa B. Fitzpatrick. **Email:**. Department of Food and Nutrition, Faculty of Agriculture and Forestry, University of Helsinki, 00014 Helsinki, Finland (Priscille Steensma). Computational Biology, Faculty of Biology, CeBiTec, Bielefeld University, 33615 Bielefeld, Germany (Marion Eisenhut). Department of Natural Product Biosynthesis, Max Planck Institute for Chemical Ecology, 07745 Jena, Germany (Maite Colinas).

## Abstract

To survive and proliferate in diverse environments with varying climate and nutrient availability, plants modulate their metabolism. Achieving a balance between carbon (C) and nitrogen (N) use such that growth and defense mechanisms can be appropriately controlled is critical for plant fitness. The identification of factors that regulate C/N utilization in plants can make a significant contribution to optimization of plant health. Here we show that pyridox(am)ine 5’-phosphate oxidase (PDX3), which regulates vitamin B_6_ homeostasis, influences C/N balance. The B_6_ vitamer imbalance resulting from loss of PDX3 leads to over-accumulation of nitrogenous compounds. A combination of increased glutamate dehydrogenase activity, impairment in the photorespiratory cycle and inappropriate use of endogenous ammonium fuel the metabolic imbalance. Growth at elevated CO_2_ levels further exacerbates the *pdx3* phenotypes. Interestingly, serine supplementation rescues growth under high CO_2_ likely bypassing the phosphorylated pathway of biosynthesis suggesting that this amino acid is an important commodity. We show that PDX3 function appears dispensable upon thermomorphogenesis, a condition that favors C metabolism. Furthermore, while a low ammonium to nitrate ratio likely accounts for overstimulation of salicylic acid (SA) defense responses in *pdx3* lines that compromises growth, a basal level of SA protects against loss of PDX3 biochemical function. Overall, the study highlights environmental scenarios where vitamin B_6_ homeostasis, as managed by the salvage pathway enzyme PDX3, is critical and provides insight into how plants reprogram their metabolism under such conditions.

## INTRODUCTION

The salvage pathways of vitamin B_6_ in plants have received increasing attention in the past years and revealed that these pathways are essential for plant development and coping with environmental stresses (Shi et al., 2002; Shi and Zhu, 2002; González et al., 2007; Herrero et al., 2011; Colinas et al., 2016; Colinas and Fitzpatrick, 2016; Gorelova et al., 2022). In plants, three enzymes have been identified so far, namely a pyridoxal (PL) kinase (SOS4) (Shi et al., 2002; Shi and Zhu, 2002), a PL reductase (PLR1) (Herrero et al., 2011) and a pyridoxine 5’-phosphate (PNP)/pyridoxamine 5’-phosphate (PMP) oxidase (PDX3) (Sang et al., 2007). Vitamin B_6_ in the form of the vitamer pyridoxal-5’-phosphate (PLP) is an important coenzyme for numerous enzymes mainly comprising amino acid biosynthesis (Liu et al., 2022). Additional functions have been attributed to vitamin B_6_ pertaining to transcription, facilitating protein folding or contributing to antioxidant activity (Liu et al., 2022). Previous studies showed that PDX3 is necessary for balancing chemical forms of vitamin B_6_ in *Arabidopsis thaliana* (referred to as Arabidopsis from here on)(Colinas et al., 2016). In the absence of PDX3 in Arabidopsis, there is accumulation of the PNP and PMP forms of vitamin B_6_ and a deficit in PLP, consistent with its biochemical function (Colinas et al., 2016). This is coincident with an alteration of metabolite levels, in particular a bias towards accumulation of nitrogenous compounds, not seen in other vitamin B_6_ metabolism mutants, and a strong morphological phenotype with reduced leaf and seed biomass (Colinas et al., 2016). A conundrum is the ability to rescue the *pdx3* phenotype by provision of nitrogen (N) in the form of exogenous ammonium (but not nitrate) despite the already increased N load in these lines (Colinas et al., 2016). Intriguingly, nitrate assimilation is impaired in *pdx3* mutants due to decreased nitrate reductase (NR) activity. Furthermore, a constitutive defense response is evident in *pdx3* lines in the absence of pathogen challenge (autoimmunity) that is coincident with an accumulation of salicylic acid (SA) (Colinas et al., 2016). Whether SA contributes to the growth defects in *pdx3* has not been explored. Moreover, the connection between vitamin B_6_ balance, N metabolism and SA-induced defense is unknown, but could provide crucial information on the importance of this vitamin for metabolic homeostasis in plants under particular environmental challenges.

A fundamental aspect of cellular homeostasis for autotrophic organisms is maintenance of carbon(C)/N balance vital for interconnecting CO_2_ fixation with N assimilation (Nunes-Nesi et al., 2010). Plants source N exogenously from their surrounding environment or endogenously through recycling of nitrogenous compounds (e.g. amino acids) or from photorespiration in the case of C_3_ plants. Under aerobic and slightly acidic conditions, the presence of nitrifying bacteria in soil leads to the oxidation of ammonium to nitrate via nitrite, thus nitrite and ammonium levels are usually low (Chalk and Smith, 2021; Subbarao and Searchinger, 2021). A considerable portion of crop cultivation of non-legumes is carried out on soil that is under such conditions and invokes the high energy consuming nitrate and nitrite reductase reactions of the plant to reduce nitrate back to ammonium for assimilation into nitrogenous compounds required by the plant for growth. Nonetheless, the paucity of N sources in current soils and the drive since decades to increase plant yields, which has greatly benefitted food security, has led to vast amounts of synthetic fertilizer being applied routinely to replenish N. However, as plants only absorb a fraction of the fertilizer applied, run-off has led to environmental pollution negatively impacting ecosystems and biodiversity (Tilman et al., 2011; Poore and Nemecek, 2018). Rising CO_2_ levels compound the problem because nitrate uptake is believed to be compromised under high CO_2_ (Bloom et al., 2010; Bloom et al., 2020). Therefore, in an effort to improve N use efficiency, knowledge of mechanisms that regulate N management by plants is critical.

In the first instance here, we explored the contribution of PDX3 to cellular vitamin B_6_ homeostasis and N management by performing metabolite analyses on Arabidopsis *pdx3* lines upon N fertilization. While ammonium application rescues the morphological phenotype of *pdx3* mutants, it masks the metabolite perturbance resulting from the incapacity of these lines to utilize soil nitrate as a source of N. Indeed, our data suggest that exogenous ammonium metabolism operates independently of PDX3. Counterintuitively, in the absence of fertilization *pdx3* mutants suffer a C/N imbalance and accumulate nitrogenous compounds (c/N). Surprisingly, exploration of photorespiration as a source of endogenous N driving this metabolic imbalance, by incubation under high CO_2_, further exacerbated the *pdx3* growth phenotype. Interestingly, the amino acid serine, which is critical for growth and N management (Zimmermann et al., 2021), alleviates the growth phenotype of *pdx3* plants under high CO_2_. Triggering of thermomorphogenesis by growth of plants at 28°C (instead of 22°C) is another condition that does not appear to require PDX3 function, and we observed that the consequent drive towards C metabolism counters the c/N imbalance in *pdx3*. Further probing of the SA-induced defense response in *pdx3* led us to unravel that is a protective strategy mediated by NPR1 and improves fitness. Overall, the study demonstrates the importance of vitamin B_6_ homeostasis as managed by the salvage pathway enzyme PDX3 to growth in diverse environments with varying nutrient availability and insight into how plants reprogram their metabolism under such conditions.

## RESULTS

### Metabolism of exogenous ammonium operates independently of *PDX3*

Our previous analyses on *pdx3* plants (*pdx3-3*/*pdx3-4*, strong and weak allele, respectively) revealed a considerable alteration in the metabolite profile compared to wild type (Colinas et al., 2016). In particular, there is a strong accumulation of nitrogenous compounds in these *pdx3* plants grown on soil in contrast to wild type. Here, in an effort to dissect the source of metabolite disturbance, we compared the metabolite profiles of soil grown *pdx3* lines and wild type plants in the absence and presence of fertilization with different N sources. In the absence of any fertilization, the profile of the *pdx3* lines was different to that of wild type with a strong enrichment of nitrogenous compounds including several amino acids and urea, corroborating previous observations (Colinas et al., 2016) (Figure 1A). There were consistent significant decreases in myo-inositol, trehalose and pyruvate, although succinate was increased (Figure 1A). In the presence of potassium nitrate fertilization, there were significant metabolite changes in *pdx3* compared to wild type, many of which were similar to those under the unfertilized conditions (Figure 1B). By contrast, supplementation with ammonium nitrate dampened the metabolite profile divergence in *pdx3* and was very similar to that of wild type under the same conditions (Figure 1C). A comparison of wild type under the unfertilized condition to that of all lines under the fertilization conditions indicated that the changes in metabolism of wild type and *pdx3* upon potassium nitrate fertilization were divergent notably in relation to certain N-rich amino acids (Figure 1D). On the other hand, the similar behavior of *pdx3* and wild type upon ammonium nitrate fertilization was particularly striking (Figure 1E). This observation suggested firstly that *pdx3* metabolizes exogenous ammonium similar to wild type and secondly the divergent metabolite profile of *pdx3* lines in the unfertilized condition is largely masked upon addition of ammonium. Notably, exogenous ammonium supply rescues the narrow, elongated, thin leaves observed with *pdx3* grown on soil, whereas nitrate does not (Supplemental Figure S1), as observed previously (Colinas et al., 2016). Nonetheless, the strong accumulation of nitrogenous compounds seen upon ammonium fertilization is what is observed for *pdx3* in the absence of fertilization, particularly the enhancement of amino acids (Figure 1A and E).

**Figure 1.**
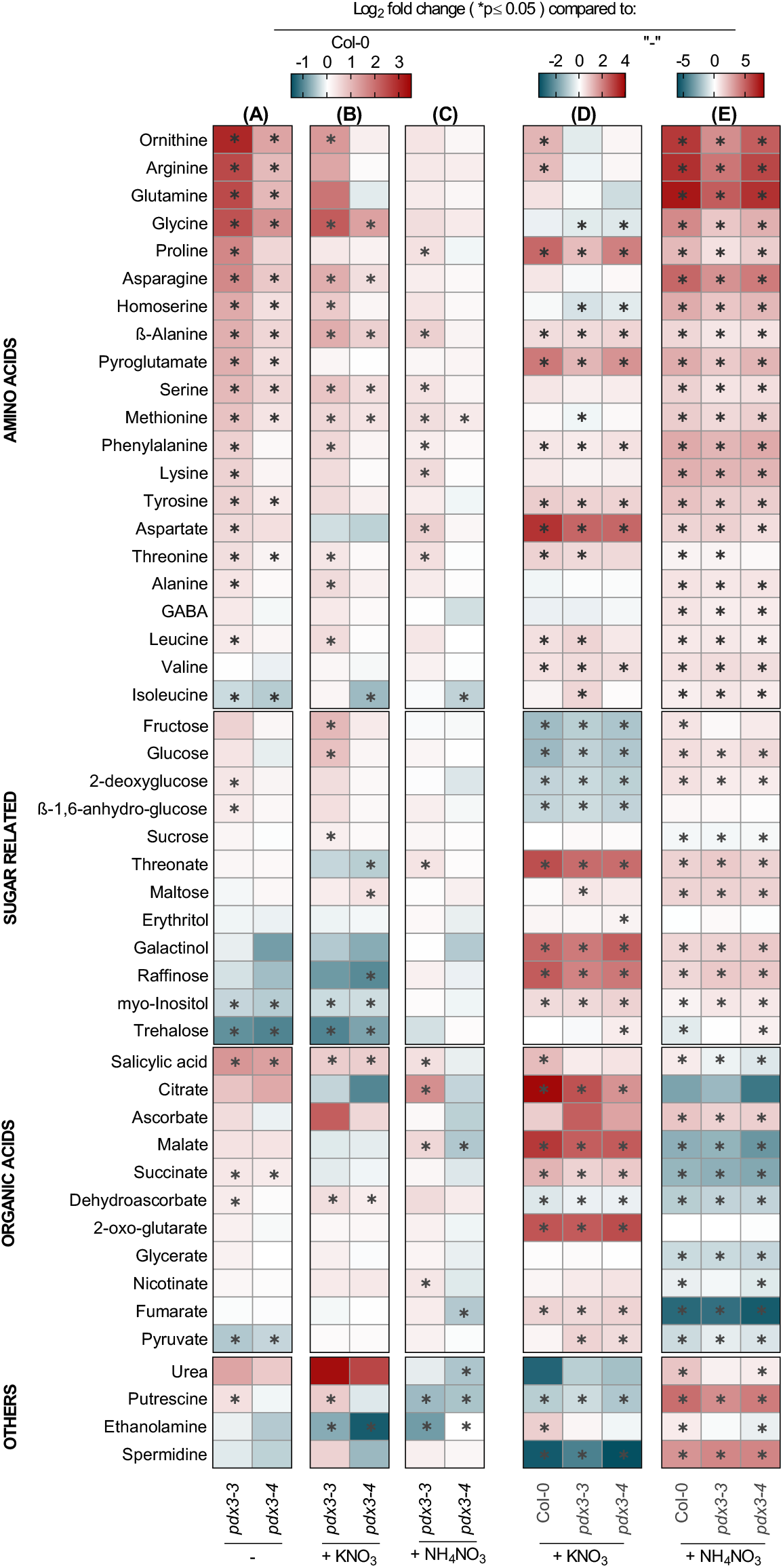
Metabolite profiles of *pdx3* plants compared to wild type under nitrogen fertilization regimes. Relative metabolite contents of *pdx3* lines are presented as heatmaps compared to wild type (Col-0) without fertilization (-) (A), potassium nitrate fertilization (+ KNO_3_) (B), or ammonium nitrate fertilization (+ NH_4_NO_3_) (C). An analysis of the same set of data in (A-C) is also presented in comparison to the metabolite profile of each respective line grown in the unfertilized condition (-). (D) compares the unfertilized condition with that of potassium nitrate fertilization (+ KNO_3_), and (E) compares the unfertilized condition with that of ammonium nitrate fertilization (+ NH_4_NO_3_). The data is represented as the Log_2_ of the average fold change (n=5-6) either to wild type (Col-0) or the condition (-). Statistical analysis was performed using a two-tailed Student’s unpaired *t*-test for p≤0.05 on fold change results using line Col-0 or condition (-) as control. The analysis was performed on 21 days old rosettes of plants grown on soil under a 16 h photoperiod (120-190 μmol photons m-2 s-1) at 22°C and 8 h darkness at 18°C. Plants were watered with water alone (-) or a 50 mM solution of the indicated compound every 9-10 days.

Given the strong bias towards enhanced N compounds in *pdx3* in the unfertilized condition, we checked molecular markers of N assimilation versus a response to exogenous ammonium via RT-qPCR. *ASPARAGINE SYNTHASE 2* (*ASN2*) is induced by exogenous ammonium (Wong et al., 2004), whereas *GLUTAMATE DEHYDROGENASE 2* (*GDH2*) induction marks ammonium assimilation (Patterson et al., 2010). Under standard growth conditions (in the absence of fertilization), *GDH2* was modestly induced in *pdx3* (particularly in the *pdx3-3* allele), whereas *ASN2* was not significantly different to wild type (Figure 2A). This was supported by mining our previous RNA-seq data from these lines grown on soil (Colinas et al., 2016), which also showed that *GDH2* was induced in *pdx3* alleles compared to wild type, whereas there was no difference in the *ASN2* levels (Figure 2A). Importantly, *GDH2* was not induced in *pdx3* lines carrying the *PDX3* transgene (complementing lines) in the absence of fertilization (Figure 2A). On the other hand, both *GDH2* and *ASN2* were induced upon ammonium nitrate fertilization in all lines but not upon potassium nitrate fertilization (Figure 2A). The aerobic conditions and slightly acidic soil used in this study is likely to contain no or very low levels of ammonium, as is also reflected by the absence of rescue of the ammonium-dependent leaf phenotype of *pdx3* mutants on unfertilized soil (Supplemental Figure S1). This lent support to a notion that there may be a C/N imbalance in *pdx3* that imparts the morphological defect. Markers of C/N imbalance include induction of *ASPARAGINE SYNTHASE 1* (*ASN1*) or *ARABIDOPSIS TOXICOS EN LEVADURA 31 (ATL31*) (Lam et al., 1994; Sato et al., 2009; Xu et al., 2019). Both were upregulated in *pdx3* compared to wild type based on mining of the previous RNA-seq data and was supported by RT-qPCR of standard soil grown samples in the absence of fertilization, notably in the *pdx3-3* allele (Figure 2B).

**Figure 2.**
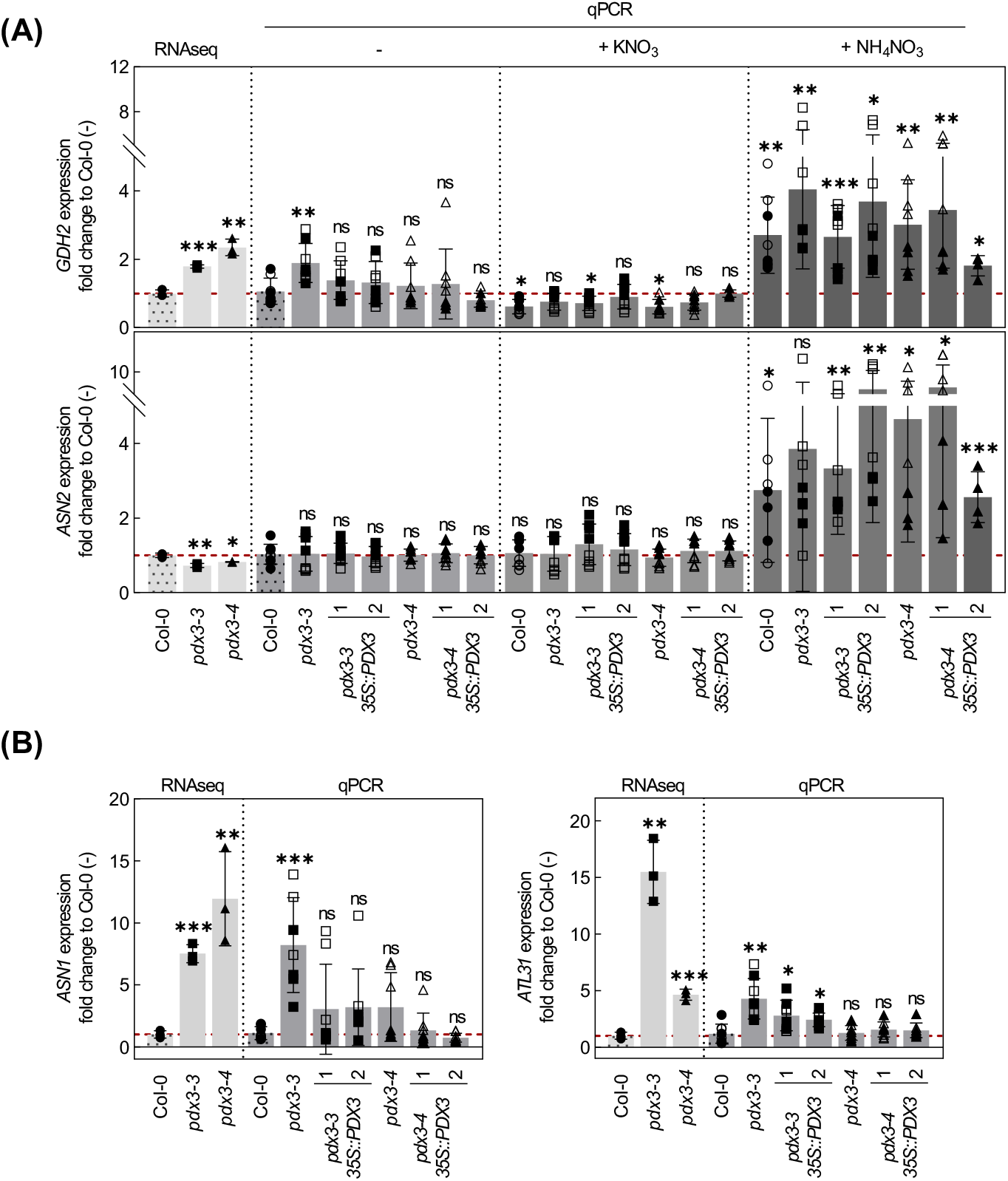
Marker gene evidence for C/N imbalance in *pdx3*. (A) Relative expression fold change, as determined by RNAseq (1) or RT-qPCR (qPCR), of glutamate dehydrogenase 2 (*GDH2*, marker for both endogenous and exogenous ammonium assimilation), and asparagine synthase 2 (*ASN2*, marker for exogenous ammonium assimilation) in wild type (Col-0), *pdx3* and complementing lines grown either on unfertilized (-), potassium nitrate (+ KNO_3_) or ammonium nitrate (+ NH_4_NO_3_) fertilized soil compared to wild type grown on unfertilized soil. (B) Asparagine synthase 1 (*ASN1*, marker for C starvation and c/N imbalance) and Arabidopsis *toxicos en levadura* 31 (*ATL31*, marker for C and N imbalance) in *pdx3* and complementing lines compared to wild type (Col-0). The data represents the mean ± SD across 2 experimental replicates (either open or filled symbols) and with 4 biological replicates each. Statistical analysis was performed using a two-tailed Student’s unpaired *t*-test with unfertilized Col-0 as control (nsp>0.05, *p≤0.05, **p≤0.005, ***p≤0.0005). The analysis was performed on 21 days old rosettes of plants grown on soil under a 16 h photoperiod (120-190 μmol photons m-2 s-1) at 22°C and 8 h darkness at 18°C. Either water alone (-) or a 50 mM solution of the indicated compound was supplemented to the soil every 9-10 days. The control was set to 1 and is indicated by the red dashed lines.

Taking all of the findings together, we infer that metabolism in the presence of exogenous ammonium operates independently of *PDX3* in Arabidopsis and thus, the requirement for *PDX3* is bypassed by ammonium fertilization. As exogenous ammonium metabolism appears to be intact in *pdx3* alleles this then masks the metabolite disturbance and consequential morphological phenotype of these mutants observed in the unfertilized soil conditions, where nitrate is the predominant source of N.

### B_6_ vitamer alteration in *pdx3* does not directly account for reduced NR activity

In contrast to ammonium fertilization, nitrate does not rescue the *pdx3* morphological phenotype as mentioned above (Supplemental Figure S1). This is not surprising because previous studies have shown that NR activity is reduced in these *pdx3* alleles (Colinas et al., 2016), although it is not known why. Biosynthesis of the NR protein is inhibited by certain amino acids as well as when ammonium is used as a source of N assimilation (Huarancca Reyes et al., 2018; Kim et al., 2018). Thus, we measured transcript levels of *NIA1/2*, the two genes coding for NR in Arabidopsis, and noted a significant reduction in *NIA1* in both *pdx3* alleles from mining of the previous RNAseq dataset, and was corroborated by RT-qPCR for the *pdx3-3* allele (Figure 3A). Protein levels of NIA1/2 were also moderately reduced in both *pdx3* alleles (Figure 3B). However, nitrate levels were unchanged compared to wild type, suggesting that nitrate uptake is intact (Figure 3C). Previously, we have also shown that the loss of PDX3 leads to an imbalance in vitamin B_6_ levels, marked by an increase in PMP and a decrease in PLP, in line with the biochemical activity of PDX3 (Colinas et al., 2016). We profiled the vitamin B_6_ content of *pdx3* alleles versus wild type upon N fertilization and observed a marked increase in PMP when ammonium was applied irrespective of the genotype as before (Colinas et al., 2016) (Figure 3D). Here, we also noted a significant (albeit modest) increase of PL in the wild type not seen in *pdx3*, and although PNP was not detectable in wild type, it was clearly observed in *pdx3* (Figure 3D). There were no other statistically significant changes in the individual vitamer profiles, but there was a general slight increase in vitamin B_6_ levels upon ammonium fertilization and no marked changes under nitrate fertilization (Figure 3D). Notably, the PMP:PLP ratio in the *pdx3* alleles remained much higher than wild type under all conditions (Figure 3E). As it is well established that NR activity decreases upon ammonium fertilization (Huarancca Reyes et al., 2018; Kim et al., 2018), we tested if PMP could directly inhibit its activity. However, PMP at concentrations up to 2.5 mM, considerably higher than found in Arabidopsis tissue (Szydlowski et al., 2013), had only a mild impact on NR activity (Supplementary Figure S2A). Thus, it is unlikely that PMP directly inhibits NR activity. Furthermore, NR activity is time of day regulated, peaking early in the day, which was maintained in *pdx3* plants, although a lower activity was observed throughout the day (Supplementary Figure S2B). We also tested if N fertilization could affect *PDX3* expression levels. However, neither transcript nor protein levels changed under either ammonium or nitrate fertilization (Supplementary Figure S2C and D). Thus, it is rather the strong accumulation of nitrogenous compounds (amino acids) in *pdx3* that is likely to trigger negative feedback regulation on NR, as reported previously for wild-type plants fed with ammonium (Huarancca Reyes et al., 2018; Kim et al., 2018). As amino acids are increased in *pdx3* even without ammonium supply and *GDH* expression is induced, we measured GDH activity and noted that it was considerably increased in the direction of glutamate formation (ammonium consumption) in the *pdx3* lines, a condition that was reversed in the complementing lines (Figure 3F). We also measured free ammonium levels and noted a significant decrease in *pdx3* compared to wild type (Figure 3G). This data suggests that internal ammonium assimilation is enhanced in *pdx3* and could at least partially account for the elevated amino acid levels. Although, decreased protein metabolism, or amino acid catabolism could also contribute.

**Figure 3.**
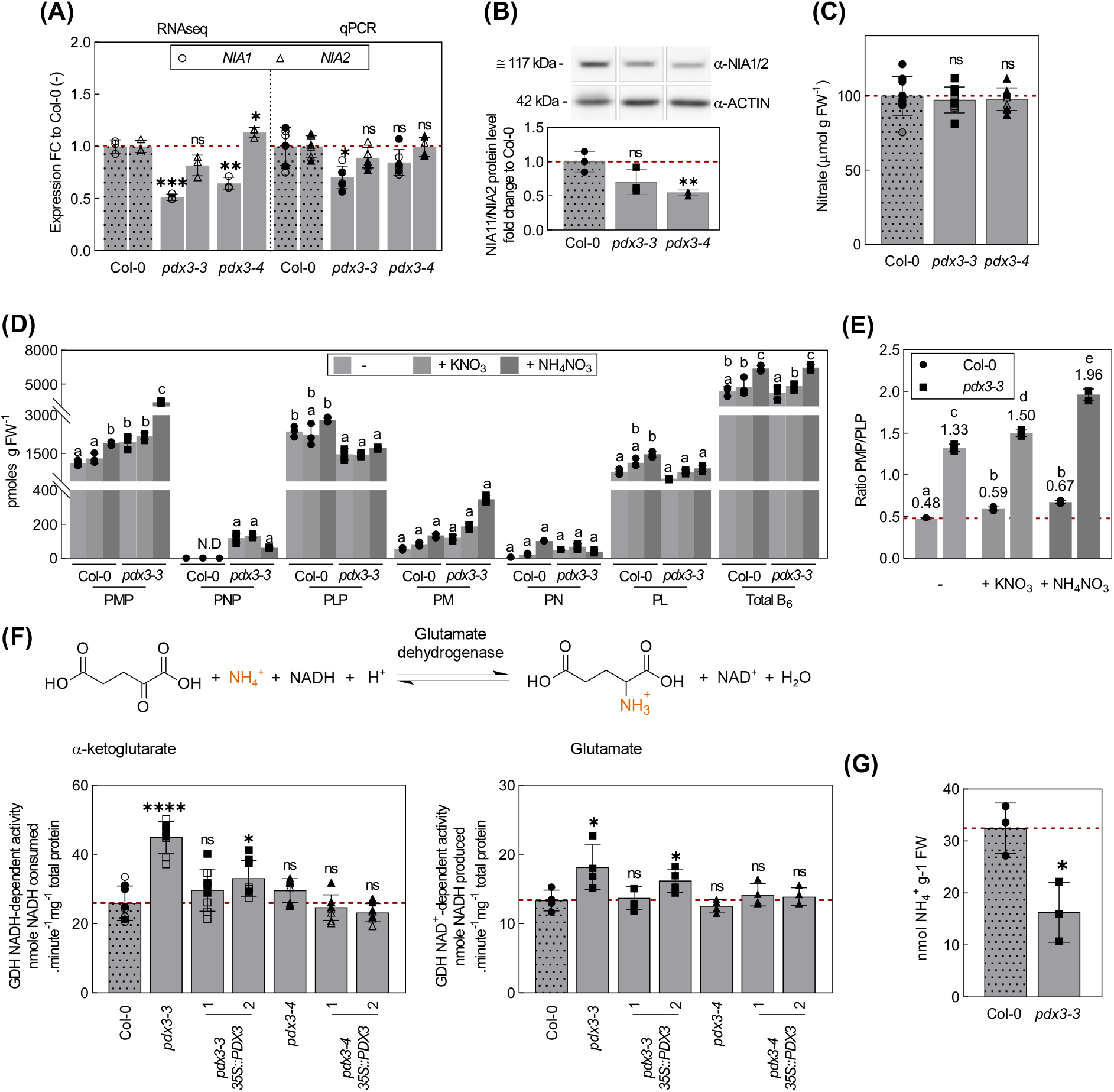
Alteration of vitamin B_6_ homeostasis does not directly impact nitrate reductase activity. (A) Relative expression fold change in expression of *NIA1* and *NIA2* in *pdx3* compared to wild type (Col-0), as determined by RNAseq (1) and by RT-qPCR (qPCR). (B) NIA1/2 protein levels of lines as in (A), determined by immunochemistry. (C) Nitrate content of lines as in (A). (D) Vitamin B_6_ profile of *pdx3-3* and wild type (Col-0) plants grown on unfertilized (-), potassium nitrate (+ KNO_3_) or ammonium nitrate (+ NH_4_NO_3_) fertilized soil. (E) PMP/PLP ratios in wild type and *pdx3-3* from the data presented in (D). (F) Glutamate dehydrogenase (GDH) activity in *pdx3* compared to wild type (Col-0). The top panel shows a scheme of the reaction; the bottom left panel shows NADH-dependent GDH activity and bottom right, NAD+-dependent GDH activity of plants. (G) Free ammonium levels in *pdx3-3* compared to wild type (Col-0). Expression data in (A and B) represent the mean ± SD across 1-2 experimental replicates (either open or filled symbols) with 3 biological replicates each. The data in (C) represent the mean ± SD across 2 experimental replicates (filled, open or semi-filled symbols) with 3-6 biological replicates each. The data in (D) and (E) represents the mean ± SD of 3 biological replicates. The data in (F) represents the mean ± SD across 1-2 experimental replicates (either open or filled symbols) with 4 biological replicates each. The data in (G) represents the mean ± SD across 2-3 biological replicates. For (A-C) and (F-G) statistical analyses were performed using a two-tailed Student’s unpaired *t*-test with unfertilized Col-0 as control (nsp>0.05 and *p≤0.05, **p≤0.005 ***p≤0.0005 and ****p≤0.00005). For (D-E) statistical analyses were performed using a two-way ANOVA with a Tukey’s multiple comparison test. Different letters indicate p≤0.05 within each vitamer sub-categories in (D), whereas in (E) different letters indicate p≤0.05 across all categories. The dashed line in (A-B) indicates the control set to 1 in (A) and (B) or the wild type (Col-0) in (C-G). The analyses in (A-F) were performed on 21 days old rosettes of plants grown on soil under a 16 h photoperiod (120- 190 μmol photons m-2 s-1) at 22°C and 8 h darkness at 18°C. Either just water (A-C, F or “-”) or a 50 mM solution of the indicated compound was supplemented to the soil every 9-10 days. The analysis in (G) was performed on 9 days old rosettes of plants grown on plates with modified MS medium containing no ammonium under a 12 h photoperiod (120 μmol photons m-2 s-1) at 22°C and 12 h darkness at 18°C.

We infer that the biochemical function of PDX3 in maintaining PMP levels does not appear to be directly related to the reduced activity of NR in *pdx3* lines, rather the accumulation of amino acids (with a contribution from enhanced ammonium assimilation) likely negatively feedback on NR biosynthesis and explain the reduced transcript/protein/activity levels.

### Growth of *pdx3* further deteriorates under high CO_2_ and is alleviated by serine supplementation

Photorespiration in C3 plants such as Arabidopsis is an important metabolic route contributing to C/N balance (Eisenhut et al., 2019). We entertained a notion that as photorespiration constrains CO_2_ assimilation and is an important source of endogenous ammonium, its repression may alleviate the C/N imbalance in *pdx3* lines and improve growth. Metabolic flux through the photorespiratory cycle can be repressed by exposing plants to elevated CO_2_ during which oxygen use by rubisco is diminished. However, exposure of *pdx3* lines to elevated CO_2_ further exacerbated the leaf phenotype of *pdx3* compared to ambient CO_2_ (Figure 4A). This suggested that contrary to our original notion, photorespiration is beneficial to *pdx3* lines. Notwithstanding, the photosynthesis rate of *pdx3* was significantly impaired under high O_2_, a condition that favors photorespiration (Figure 4B).

**Figure 4.**
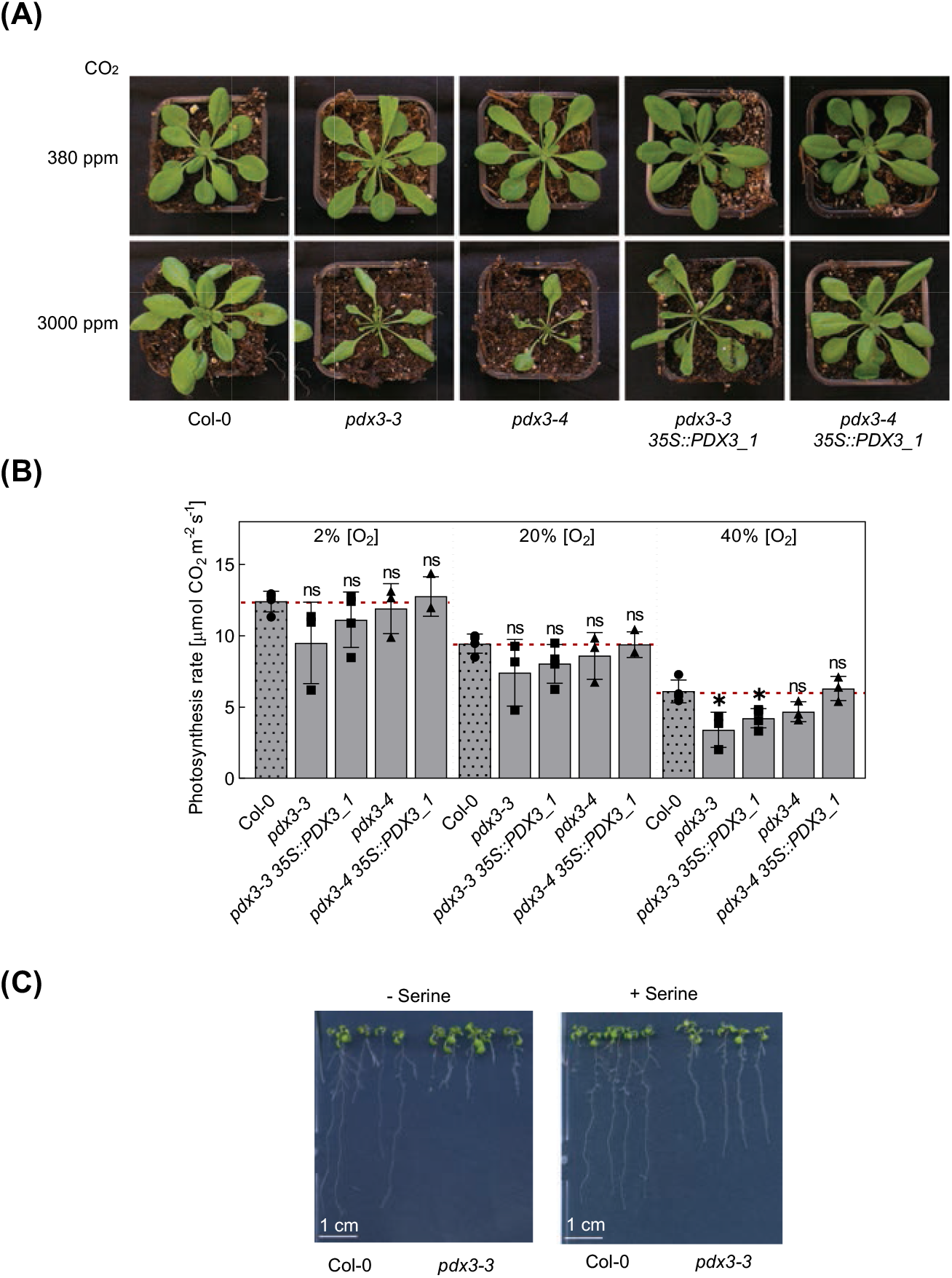
Growth of *pdx3* further deteriorates under high CO_2_ and is alleviated by serine supplementation. (A) Phenotypic comparison of wild type (Col-0), *pdx3* and complementing lines grown under elevated CO_2_ (3000 ppm) or ambient CO_2_ (380 ppm) on soil and a 12 h photoperiod (120 μmol photons.m^−2^.s^−1^) at 22°C and 12 h of darkness at 18°C. (B) Analysis of the rate of photosynthesis as a function of the oxygen percentage in *pdx3* and complementing lines compared to wild type (Col-0). Plants were grown under the same conditions as in (A). The data represents the mean ±SD of 3 experimental replicates. Statistical analysis was performed using a multiple two-tailed Student’s unpaired *t*-test using the Holm-Sidak method with Col-0 as control (^ns^p>0.05 and *p<0.05). (C) Growth of wild type (Col-0) and *pdx3* on culture plates under elevated CO_2_ (3000 ppm) in the absence or presence of serine (100 μM) supplemented to the medium.

We remained puzzled by the bias towards enhanced nitrogenous compounds in *pdx3* and strived to define which cellular functions were perturbed by the lack of PDX3 activity. Recently, the phosphorylated pathway of serine biosynthesis (PPSB) has been shown to be vital for Arabidopsis plant growth (Zimmermann et al., 2021). The PPSB pathway is predominantly active in plant roots and is a major contributor to the serine pool, especially under elevated CO_2_, when photorespiration is repressed (Zimmermann et al., 2021). Mutations in the PPSB pathway lead to an accumulation of nitrogenous compounds and a severe impact on growth, particularly under high CO_2_ conditions (Zimmermann et al., 2021). The parallels with the phenotypes of *pdx3* allowed us to postulate that *pdx3* plants may suffer a deficiency in serine biosynthesis through the PPSB pathway. This notion was supported by the fact that 3-phosphoserine aminotransferase (PSAT), an enzyme of the PPSB pathway, is dependent on PLP for activity (Wulfert and Krueger, 2018). As external feeding with serine has been shown to be metabolized the same way as the plant would by biosynthesis *de novo* (Zimmermann et al., 2021), we tested if serine supplementation could improve growth of *pdx3* plants under elevated CO_2_. Indeed, growth of *pdx3* on culture plates under elevated CO_2_ was considerably improved in the presence of serine (Figure 4C).

This suggests that serine biosynthesis through the PPSB pathway is compromised in *pdx3* plants and contributes to the C/N imbalance in these lines. Moreover, our data clearly indicate that PDX3 function is crucial under elevated CO_2_ levels.

### A shift in metabolism at elevated temperatures compensates for loss of *PDX3*

One other feature of *pdx3* lines is that they are morphologically similar to wild type plants when grown at 28ºC (Colinas and Fitzpatrick, 2016). In an effort to further probe the C/N imbalance in *pdx3*, we determined the metabolic status of lines grown at the standard 22ºC to that at 28ºC. As Arabidopsis plants proceed through developmental transitions faster at higher temperatures, we studied rosette leaves at either 14 or 12 days after germination at 22°C and 28°C, respectively, timepoints at which five true leaves were present under both conditions (Supplemental Figure 3). At 28°C there was a clear shift in the metabolic profile compared to 22°C such that the proportion of sugar-related compounds was higher in wild type as well as in the *pdx3* lines (Figure 5A and B). Thus, there was a reduction in the abundance of nitrogenous compounds in *pdx3* at the higher temperature and the proportional distribution of metabolites was largely similar to wild type at 28°C. However, the vitamin B_6_ profile of *pdx3* mutant lines showed largely the same perturbations compared to wild type at both temperatures (Figure 5C). Notably, the vitamin B_6_ profile of wild type was not substantially altered at 28°C compared to 22°C (Figure 5C).

**Figure 5.**
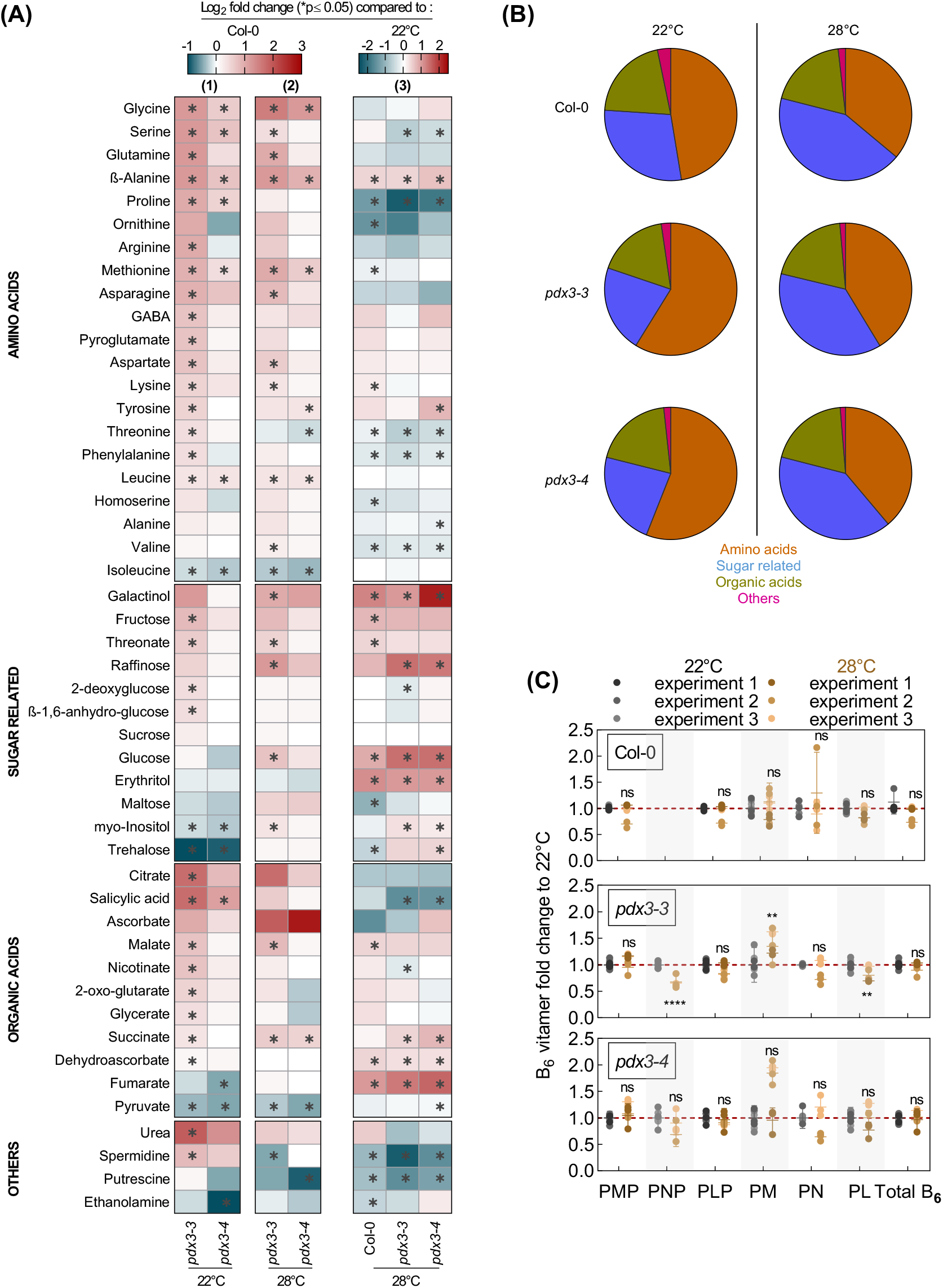
Metabolite profiling of wild type and *pdx3* plants grown under different temperatures. (A) Relative metabolite contents of *pdx3* lines are presented as heatmaps compared to wild type (Col-0) at 22°C (panel 1), or of *pdx3* at 28°C compared to 22°C (panel 2), or of wild type and *pdx3* compared to wild type at 22°C (panel 3). The data is represented as the Log_2_ of the average fold change (n=5-6) to Col-0 or condition (22°C). Statistical analysis was performed using a two-tailed Student’s unpaired *t*-test for p≤0.05 on fold change results using line Col-0 or condition (22°C) as control. (B) Pie-chart representation of the proportions of relative abundance of each group of metabolites within the total amount of metabolites measured. (C) Fold change in B_6_ vitamers of wild type and *pdx3* lines grown at 28°C (brown) compared to 22°C (gray). The data represents the mean ± SD across three independent experimental replicates (represented by different shades of brown and gray, respectively). Statistical analysis was performed using a two-tailed Student’s unpaired *t*-test using the individual vitamers at 22°C as control (nsp>0.05, **p≤0.05 and ****p≤0.0005). The control (vitamer at 22°C) was set to 1 and is indicated by the red dashed line. The analysis was performed on rosettes of plants grown on soil (unfertilized) up to 14 days after germination under a 16 h photoperiod (120-190 μmol photons m-2 s-1) at 22°C and 8 h darkness at 18°C (22°C) or up to 12 days after germination at constant 28°C (28°C).

The data indicate that there is a general shift in C/N balance at 28°C with a higher proportion of C compounds and less investment in N compounds compared to that at 22°C. The metabolic state at elevated temperature likely compensates the C/N imbalance observed in *pdx3* at 22°C and thus growth of *pdx3* matches that of wild type at 28°C. These observations suggest metabolism at 28°C is not reliant on PDX3 and interestingly is not noticeably affected by the imbalance of B_6_ vitamers, in contrast to 22°C.

### Molecular manipulation to decrease elevated SA levels in *pdx3* improves growth

Another prominent feature of *pdx3* lines grown under standard conditions on soil, is the elevation of transcripts related to the defense hormone SA, as well as SA itself (Colinas et al., 2016; Colinas and Fitzpatrick, 2016). A central dogma in plant science is that growth is compromised when plants are in defense mode fending off attack (He et al., 2022). Interestingly, this trade-off can be alleviated by elevated temperatures and ammonium fertilization both of which reduce SA levels (Wang et al., 2013; Kim et al., 2021). In this context, we observed here that the SA content was considerably reduced in *pdx3* at 28°C compared to 22°C, as well as by fertilization with ammonium (Figure 6A), i.e. conditions in which *pdx3* behaves like wild type. To dissect the contribution of SA to the phenotype of *pdx3* at 22°C on unfertilized soil, we used a genetic approach to reduce SA levels. In particular, we crossed *pdx3* lines with the *SALICYLIC ACID DEFICIENT 2* mutant line (*sid2-1*) in which the isochorismate pathway of SA biosynthesis is blocked (Nawrath and Metraux, 1999), as well as with the transgenic line carrying the *Pseudomonas putida NahG* gene that encodes salicylate hydroxylase and degrades SA into catechol (Lawton et al., 1995). While monitoring the growth of plants on soil throughout the vegetative stage of development, we noted that the leaf morphological defect in *pdx3* lines was already visible with the development of the first true leaves, which were narrower, malformed in shape and had a reduced leaf lamina area compared to wild type (Figure 6B). This feature was characteristic of all newly emerging leaves (Figure 6B). On the other hand, these deformities were alleviated somewhat in the *pdx3-3 sid2-1, pdx3-4 sid2-1* or *pdx3-3 NahG* or *pdx3-4 NahG* double mutant lines (Figure 6B). In line with this, the expression of *PATHOGEN RESISTANT1* (*PR1*) was drastically reduced in these lines (Figure 6C). However, we noted that although the leaf phenotype was improved, deformities in shape were still evident in developing leaves and lesions were present that are not found in *pdx3* alone (Figure 6B). Thus, while diminishing SA levels through genetic manipulation improves *pdx3-3* and *pdx3-4* growth, it does not fully support a hypothesis where growth is compromised solely because of an SA-triggered defense response in *pdx3-3* and *pdx3-4* mutants. Interestingly, the increased transcript levels of *ASN1, ATL31, GDH2* seen in *pdx3* alone, as markers of C/N imbalance, approached those of wild type in the *pdx3-3 NahG* and *pdx3-3 sid2-1* double mutants (Figure 6C). Of note, these transcript levels were similar between wild type and the single *NahG* and *sid2-1* mutants at 22°C (Figure 6C). This suggested that SA likely contributes to C/N imbalance in *pdx3*. To probe this finding further, we crossed *pdx3-3* with *nonexpressor of PR genes 1-2* (*npr1-2*) (Cao et al., 1997) that is unable to transduce SA defense responses. Interestingly, the *pdx3-3 npr1-2* double mutant was not effective in alleviating the developmental defects of *pdx3-3*, in contrast to the *pdx3-3 sid2-1* or *pdx3-3 nahG* double mutant lines, indeed it appeared even worse (Figure 6B). In this context, it should be realized that SA can still be biosynthesized through alternative pathways, e.g. via phenylalanine ammonia lyase and transduced (requiring NPR1) in *pdx3-3 sid2-1* and *pdx3-3 nahG* mutant lines as well as the individual *sid2-1* and *NahG* mutants (Nair et al., 2021). This implies that a basal level of SA and the corresponding signaling (via NPR1) benefits *pdx3* growth. Notably, the individual *sid2-1, nahG*, and *npr1-2* lines developed similar to wild type plants under our conditions (Figure 6B).

**Figure 6.**
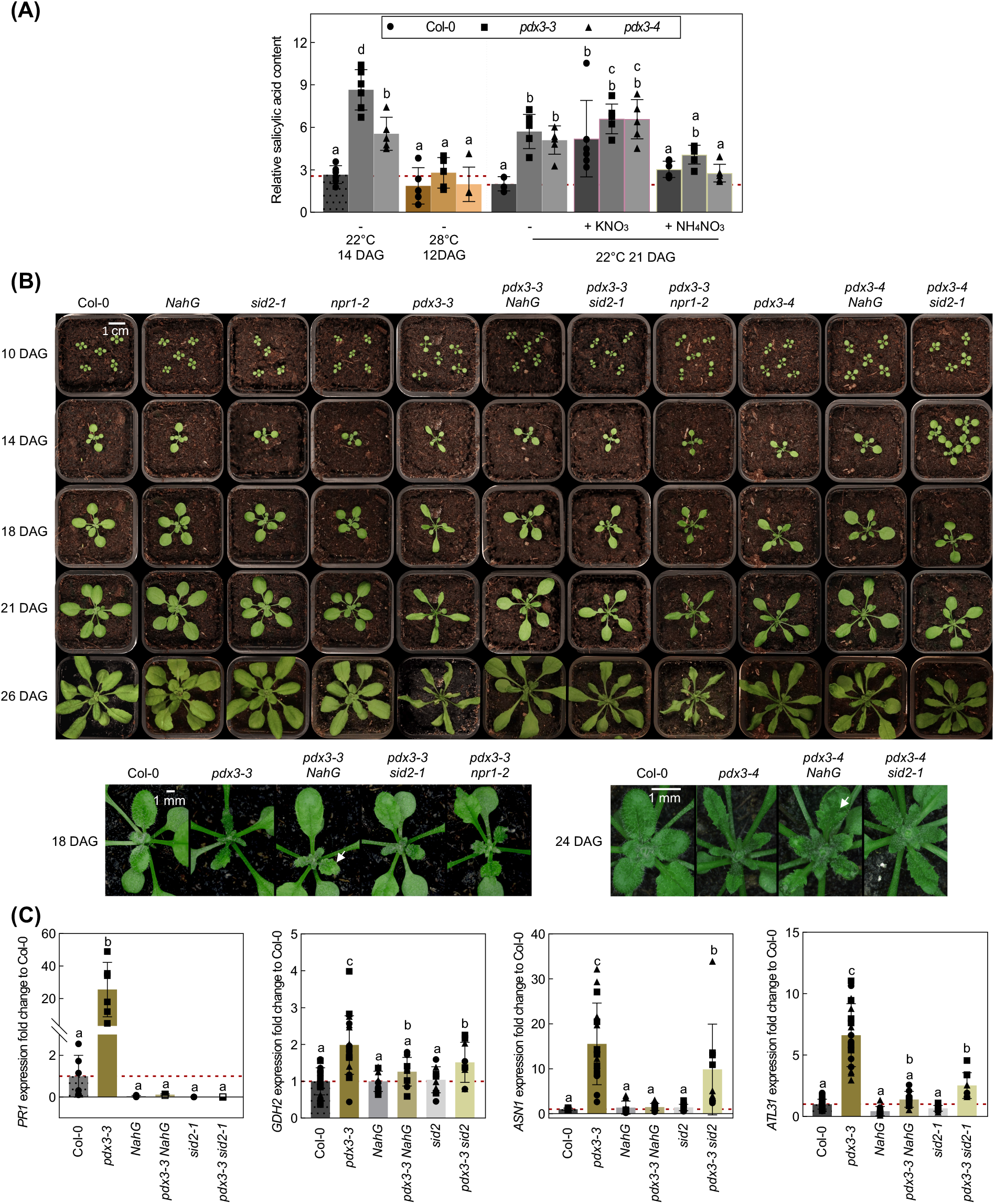
Salicylic acid (SA) contributes significantly to the *pdx3* leaf phenotype. (A) Relative SA content in 12, 14, and 21 days old wild type (Col-0) and *pdx3* lines as indicated. Plants were grown on soil under either a 16 h photoperiod (120-190 μmol photons m-2 s-1) at 22°C and 8 h darkness at 18°C (22°C) or a 16 h photoperiod at 28°C and 8 h darkness at 28°C (28°C). Either water alone (-) or a 50 mM solution of the indicated compound was supplemented to the soil every 9-10 days. The data represents the mean ± SD of 5-6 biological replicates. (B) Photographs of wild type (Col-0), *sid2-1, NahG, npr1-2, pdx3* lines, and *pdx3-3 sid2-1, pdx3-3 NahG, pdx3-3 npr1-2, pdx3-4 sid2-1, and pdx3-4 NahG* at various days after germination (DAG) as indicated. The lower left panel is a close-up of the corresponding lines shown in the upper panel. The lower right panel are close-up photos of the lines as indicated 24 DAG. White arrows indicate leaf lesions. (C) Relative expression fold change, as determined by RT-qPCR, of *PATHOGENESIS RELATED PROTEIN1* (*PR1*), *Glutamate dehydrogenase 2* (GDH2), *Asparagine synthase 1* (*ASN1*), and *Arabidopsis toxicos en levadura 31* (ATL31) in 21 days old wild type (Col-0), *pdx3-3, NahG, pdx3-3 NahG, sid2-1*, and *pdx3-3 sid2-1*. The data represents the mean ± SD across three independent experimental replicates (different squares, dots and triangles) of 3-6 biological replicates each. Statistical analysis in (A) and (C) was performed using a one-way ANOVA with a Fisher’s LSD test (different letters indicate p≤0.05). The level of the control (Col-0) in (A) or the control set to 1 (C) is indicated by the red dashed line. The analysis in (C) was performed on rosette leaves of soil grown plants (unfertilized) under a 16 h photoperiod (120-190 μmol photons m-2 s-1) at 22°C and 8 h darkness at 18°C.

Taken together the data suggests that while SA overaccumulation partially contributes to the developmental and morphological defects in *pdx3* mutants, it is not the primary cause of these abnormalities. Moreover, a basal level of SA that potentiates signaling (via NPR1) prevents even poorer performance of *pdx3*.

### SA modulates vitamin B_6_ metabolism but cannot overcome a lack of PDX3 function

We next examined for a possible connection between SA and vitamin B_6_ homeostasis applicable to the loss of PDX3. In the first instance, we determined the vitamin B_6_ profile of *sid2-1* and *NahG* compared to wild type. We noted that there was a significant decrease in PMP, PM and PL that drove an overall decrease in vitamin B_6_ levels in *NahG* in particular compared to wild type (Figure 7A). Noteworthy, this was also sometimes observed in *sid2-1* but was inconsistent. On the other hand, the vitamin B_6_ profile of *pdx3-3 sid2-1* and *pdx3-3 NahG* mutant lines was largely similar to that of *pdx3* alone (Figure 7B). To test if SA may contribute to modulation of vitamin B_6_ contents, we treated wild type plants with SA and determined the vitamin B_6_ profile. We observed an increase in PM in particular, as well as PL, albeit more modest (Figure 7C). The increase was transient and had reverted back to the original levels 48 hours after treatment, coincident with a visible recovery of the plants from the treatment and repression of the induction of *PR1* (Figure 7D). To test the importance of the transient response in wild type, we also determined the vitamin B_6_ profile of Arabidopsis lines that constitutively accumulate SA and, in this context, phenocopying *pdx3*. In particular, we used the *BONZAI1* mutant (*bon1-1)* and the gain of function *SUPRESSOR OF npr1-1, CONSTITUTIVE 1* mutant (*snc1-1)* (Hua et al., 2001; Li et al., 2001; Zhang et al., 2003; Yang and Hua, 2004). *BON1* is a regulator of growth and defense homeostasis that mediates its response by negatively regulating the haplotype specific *R* gene, *SNC1*. The *bon1-1* line had significantly increased levels of PLP (as well as PM and PL but not always consistent) that drove a modest increase in total vitamin B_6_ levels (Figure 7E). Interestingly, *snc1-1* had significantly increased levels of PMP, PLP and PL that resulted in an increase in total vitamin B_6_ levels (Figure 7E).

**Figure 7.**
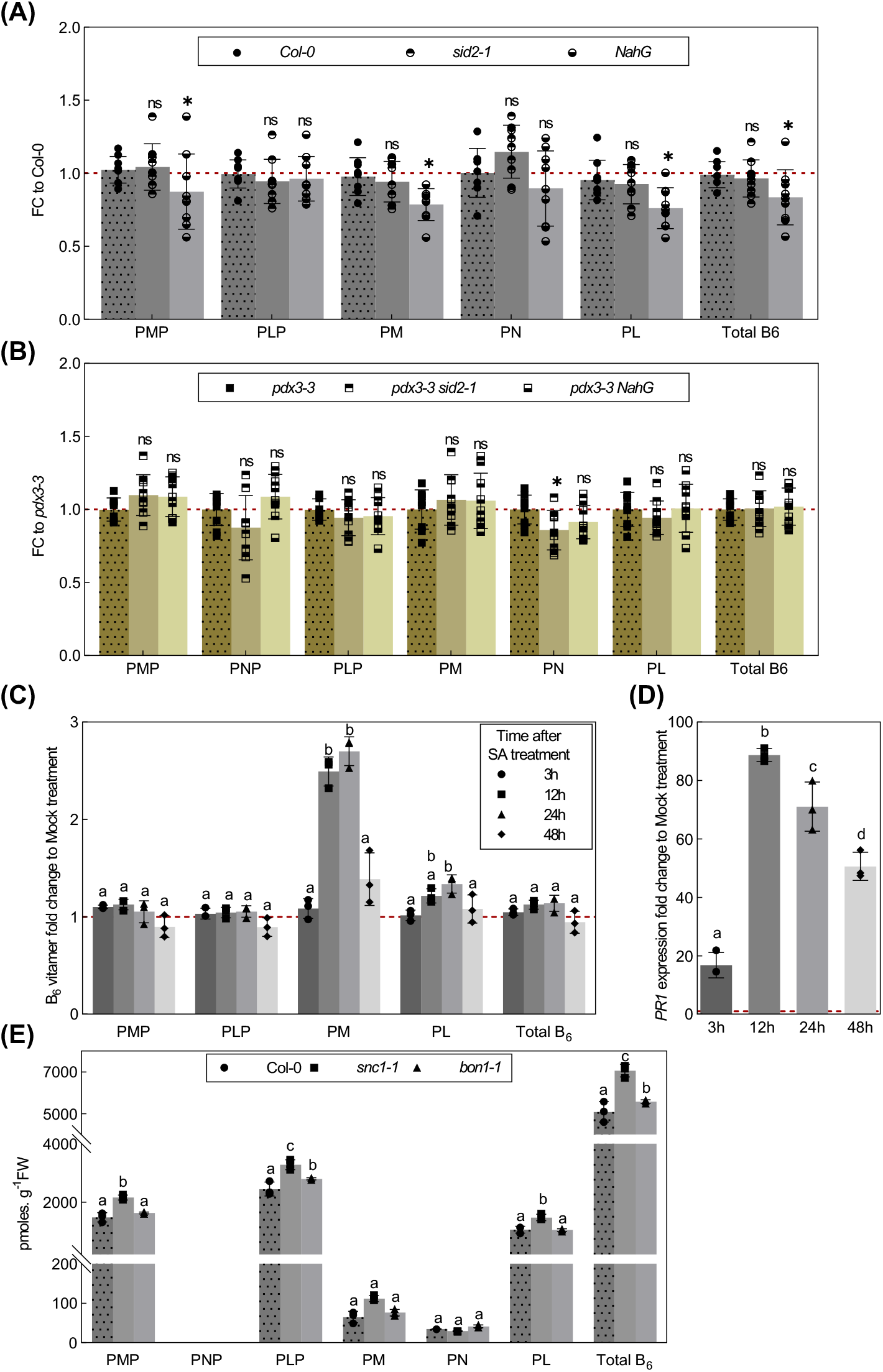
Salicylic acid can alter vitamin B_6_ levels. (A) B_6_ vitamer content fold change (FC) of 21 days old *sid2-1* and *NahG* plants normalized to the B_6_ vitamer content of wild type (Col-0) or (B) of *pdx3-3 sid2-1* and *pdx3-3 NahG* normalized to the B_6_ vitamer content in *pdx3-3* alone. (C) Fold change of the vitamin B_6_ profile of salicylic acid (SA) treated 19 days old wild type (Col-0) at 0 h, 3 h or 12 h after treatment, as well as 20 days old and 21 days-old, at 24 h and 48 h after treatment, respectively where each time point was normalized to the corresponding mock treated plant. (D) Relative expression fold change (RT-qPCR) of *PATHOGENESIS RELATED PROTEIN1* (*PR1*) of SA-treated plants from (C) at each time point compared to the respective mock treatment. (E) Vitamin B_6_ profile and cumulative total vitamin B_6_ of 21 days old wild type (Col-0) and the autoimmune mutants *snc1-1* (gain of function point mutation that leads to a constitutively active protein) and *bon1-1* (loss of function). The data in (A-B) represents the mean ±SD across 3 independent experimental replicates of 3 biological replicates each. The data in (C-E) represents the mean ±SD of 3 biological replicates. Statistical analysis in (A-B) was performed using a two-tailed Student’s unpaired *t*-test using Col-0 or *pdx3-3* as control (nsp>0.05, *p≤0.005 and ***p≤0.005). Statistical analysis in (C) and (E) was performed using a two-way ANOVA with Tukey’s multiple comparisons test (different letters indicate p≤0.05). Statistical analysis in (D) was performed using one-way ANOVA with Tukey’s multiple comparisons test (different letters indicate p≤0.05). The analyses were performed on rosettes of plants grown on soil (unfertilized) under a 16 h photoperiod (120-190 μmol photons m-2 s-1) at 22°C and 8 h darkness at 18°C.

Taken together, we infer that SA alters vitamin B_6_ metabolism. These effects do not mimic those of *pdx3* but may (partially) compensate the vitamin B_6_ imbalance and thus *pdx3* plants fare better with SA-signaling dependent responses. Nonetheless, Arabidopsis plants that lack PDX3 cannot completely overcome the vitamin B_6_ imbalance (increased PMP:PLP) that drives a c/N imbalance and as a consequence overaccumulation of SA at 22°C on soil, when nitrate is the predominant source of N. Ammonium fertilization or elevated temperatures (at ambient CO_2_) that drive a different metabolism (coincident with reducing SA levels) bypass the requirement for the biochemical function of PDX3.

## DISCUSSION

Here we build on previous work (Colinas et al., 2016) to highlight the importance of the salvage pathway enzyme PDX3 in maintaining vitamin B_6_ balance for metabolism under distinct environmental conditions. Under standard ambient conditions for Arabidopsis, the absence of PDX3 compromises C/N balance through accumulation of nitrogenous compounds, and SA-related defenses are triggered. The metabolic imbalance and its negative impact on growth of *pdx3* lines can be fully alleviated by ammonium fertilization or growth of plants under higher temperatures, which as we show here do not appear to rely on the biochemical function of PDX3 due to a different metabolic homeostasis under these conditions. These conditions also dampen the overstimulated SA related defense response in *pdx3* and can be partially mimicked by crossing lines diminished in SA content with *pdx3*. However, repression of SA signaling further compromises *pdx3* growth, suggesting *pdx3* co-opts SA for increased fitness. Intriguingly, our data also show that PDX3 function is particularly important under elevated CO_2_ conditions – a feature associated with serine biosynthesis, thus denoting a distinct function for this protein in its contribution to vitamin B_6_ and C/N balance.

We first examined the consequences of N fertilization on the metabolite profile of *pdx3* lines. The suppression of the morphological phenotype of *pdx3* by supplementation with ammonium could be explained as bypassing a nutritional deficiency in *pdx3* when nitrate is the source of N. However, this simple explanation is not sufficient to describe observations in this study and suggest a more complex role for PDX3 in managing endogenous N metabolism. Firstly, the metabolite profile of *pdx3* leaves is largely indistinguishable from that of wild type upon ammonium fertilization, which suggests that metabolism of exogenous ammonium is intact in *pdx3*. Notably, assimilation of exogenous ammonium is diagnosed by accumulation of several nitrogenous compounds, e.g. amino acids, reduction of NR activity, as well as an increase in the level of certain B_6_ vitamers such as both PMP and PLP. Some of these features are also diagnostic of the absence of PDX3. However, in *pdx3* mutants, PMP increases due to the lack of PMP oxidase activity and there is a deficit in PLP, and thus the PMP:PLP ratio is not maintained (see scheme in Figure 8). During transaminase reactions that allow the interconversion of amino and keto acids (Koper et al., 2022), PMP is a natural intermediate derived from the coenzyme PLP, that facilitates the transfer of an amino group to a keto acid to make an amino acid (Figure 8). An increase in PMP levels (as seen in *pdx3*) may drive the equilibrium in favor of amino acid formation (which we refer to as the “N-gear”, Figure 8). Indeed, the observed decreased levels of free ammonium (Figure 3G) support the notion of increased N assimilation in *pdx3*, albeit from endogenous sources (see below). Therefore, we propose that *pdx3* plants suffer a C/N imbalance (Figure 8), which is supported by the induction of marker genes *ASN1* and *ATL31*. As C skeletons are required also for biosynthesis of nitrogenous compounds, a shift to the “N-gear” in *pdx3* would compromise C oxidation for energy generation and would account for the impaired growth of the plants, an association that can be investigated in future studies. Notably, C limitation may also trigger a higher activity of SNF1-related kinase1 (SnRK1) (Mair et al., 2015), which in *pdx3* could also explain the observed decreases in protein levels of NR, as its phosphorylation by SnRK1 allows for the anchoring of 14.3.3 proteins that in turn are targets for ubiquitination by ubiquitin ligases such as ATL31 (Lillo et al., 2004; Polge et al., 2008; Huarancca Reyes et al., 2018). Notwithstanding, the accumulation of amino acids is known to negatively regulate biosynthesis of NR (Huarancca Reyes et al., 2018). Nitrate levels are similar to wild type in *pdx3*, suggesting that nitrate uptake is not impacted in this mutant. The notion that PDX3 is not required for exogenous ammonium assimilation is supported by the recent implication of other vitamin B_6_ genes in this process. Specifically, the biosynthesis *de novo* gene *PDX1*.*1* is transcriptionally induced in Arabidopsis seedlings grown on medium supplemented with ammonium (Liu et al., 2022) and ascribed to the associated management of reactive oxygen species (Swift et al., 2020; Liu et al., 2022). Further, the kinase that phosphorylates B_6_ vitamers, SOS4, has been implicated in management of nitric oxide levels that also accompanies exogenous ammonium uptake (Zhang et al., 2021). Such mechanisms under ammonium nutrition would mask the negative effects seen when *PDX3* is absent under nitrate nutrition or on unfertilized soil (where nitrate is the predominant source of N) and thus management of exogenous ammonium likely operates independently of PDX3.

**Figure 8.**
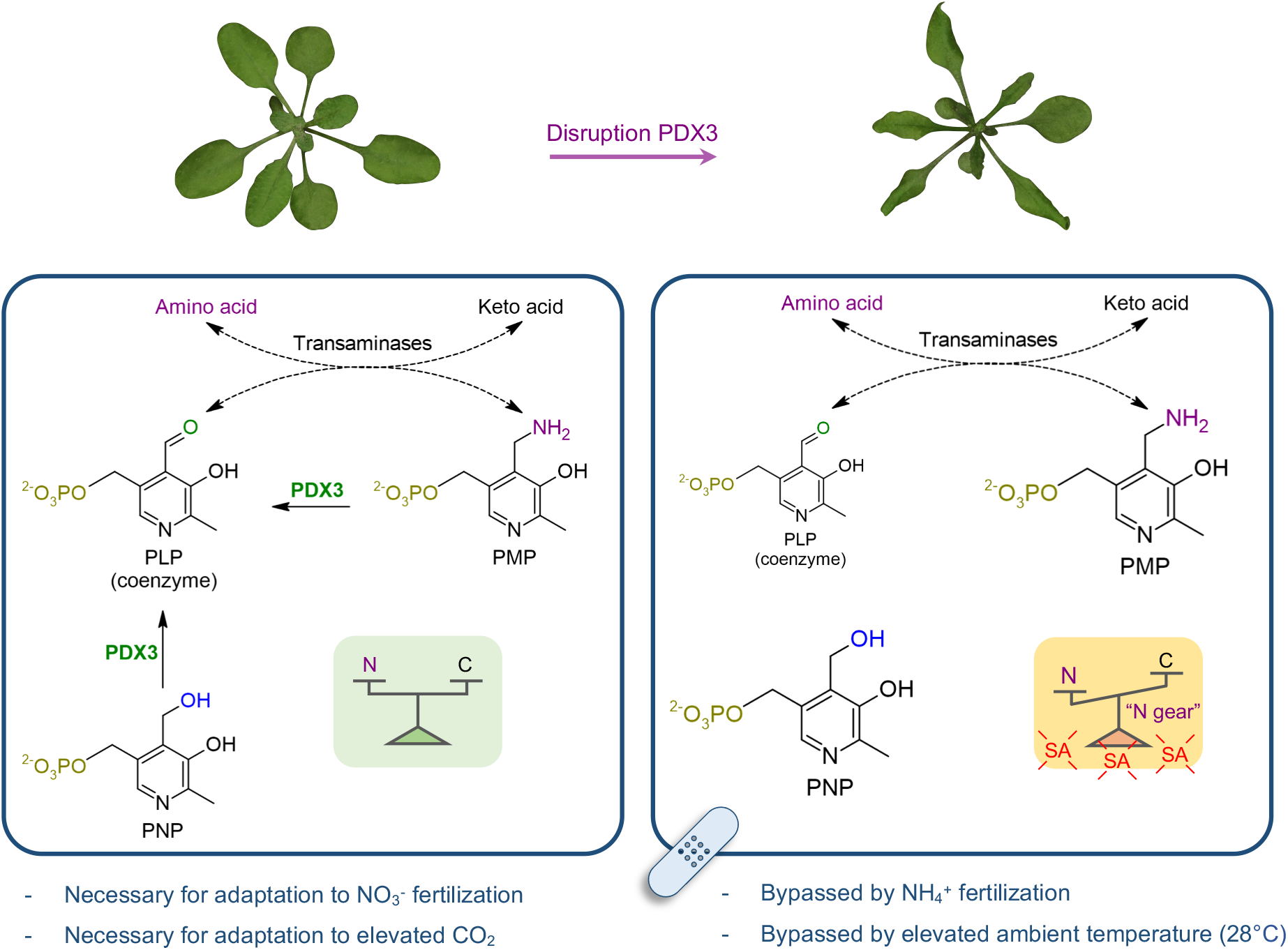
Impact of PDX3 on C/N balance in Arabidopsis. Left panel: PDX3 serves to balance the B_6_ vitamers pyridoxamine 5’-phosphate (PMP) and pyridoxal 5’-phosphate (PLP) both of which are coenzyme intermediates during amino acid and keto acid interconversions by transaminases. This in turn contributes to carbon (C) and nitrogen (N) balance optimal for plant growth, particularly under nitrate (NO -) fertilization and elevated CO_2_ conditions. PDX3 can also use pyridoxine 5’-phosphate PNP to form PLP. Right panel: In the absence of PDX3, PMP and PNP accumulate and there is less PLP that perturbs the C/N equilibrium in favor of N (“N-gear”). This is accompanied by a salicylic acid (SA) defense response and negatively impacts plant growth and fitness. The morphological phenotype resulting from a lack of PDX3 function can be bypassed (band-aid) by ammonium (NH_4_+) fertilization or elevated ambient temperatures (28°C).

As nitrate assimilation is impaired in *pdx3*, we infer that N is sourced internally to furnish accumulation of nitrogenous compounds in *pdx3* when grown on soil. N as ammonium can be released and recycled in the plant through photorespiration and the degradation of N-compounds such as arginine (Coruzzi, 2003; Hildebrandt et al., 2015). Indeed, the accumulation of urea and ornithine in *pdx3* indicates arginine degradation, even though steady state arginine levels are high. Of note also is an early study reporting inhibition of GDH by PLP and inversely promotion of activity in the direction of ammonium release and assimilation into amino acids when PLP levels are low (Teixeira and Davies, 1974). The lower levels of PLP in *pdx3* could account for the corresponding increase in GDH deaminating activity observed here and contribute to the increase in amino acids. Ammonium release from photorespiration could also fuel amino acid biosynthesis but the exaggeration of the *pdx3* phenotype under high CO_2_ when photorespiration is diminished (Benstein et al., 2013) allows us to deduce that impaired photorespiration is not the primary cause of the growth phenotype. Although, as nitrate assimilation is reduced under high CO_2_ (Bloom et al., 2010; Bloom, 2015; Bloom et al., 2020), this could account for the more drastic phenotype under this condition. This is in contrast to the recently described *er-ant1* mutant which suffers a PLP deficiency that impacts photorespiration and is alleviated by growth under high CO_2_ (Altensell et al., 2022). Thus, the *pdx3* phenotype is a consequence of a different metabolic defect. These observations led us to serine biosynthesis and the importance of this amino acid for growth and N metabolism as recently illustrated by the Krueger group (Wulfert and Krueger, 2018; Zimmermann et al., 2021). Interestingly, impairment of the PPSB pathway triggers an alteration in N metabolism, exemplified by enhanced ammonium assimilation and an increase in amino acids, as also observed here for *pdx3* (Zimmermann et al., 2021). The contribution of PDX3 to this process is supported by the rescue of *pdx3* growth upon exogenous serine supply, particularly under high CO_2_ conditions, when serine biosynthesis through photorespiration in Arabidopsis is compromised and the PPSB route that relies on PLP is crucial. Notably, PLP is also a key coenzyme for glycine decarboxylase (P-protein) and moreover PNP has been reported to inhibit its functionality in bacteria (Ito et al., 2019). The deficit in PLP and increase in PNP observed in *pdx3* may thus also limit serine supply through photorespiration which is exasperated under high CO_2_.

Compromising PDX3 biochemical function renders the plant with an autoimmune phenotype as reported previously (Colinas et al., 2016), that reduces growth under the standard temperature of 22°C. Previous studies on SMALL UBIQUITIN-RELATED MODIFIER (SUMO), *siz1-2* and *siz1-3* and *nudt6-2 nudt7* have shown that a low ammonium to nitrate ratio triggers SA accumulation and induction of marker transcripts, such as *PR1* that can be alleviated by ammonium fertilization (Park et al., 2011; Wang et al., 2013; Kim et al., 2021). A similar mechanism may operate in *pdx3* that compromises growth, as the ammonium content is decreased and nitrate content is similar to wild type under standard growth conditions (22°C) without fertilization. On the other hand, the wild type behavior of *pdx3* lines at 28°C can be explained by triggering of the thermomorphogenesis program mediated by PIF4, which alongside SIZ1-dependent SUMOylation facilitates the downregulation of SNC1-dependent immunity and SA biosynthesis, to favor growth (Gangappa et al., 2017; Hammoudi et al., 2018). In accordance with this, defense gene transcripts and SA were reduced in *pdx3* at 28°C. Initially, we then inferred that the morphological phenotype of *pdx3* is purely a consequence of SA-triggered autoimmunity, that may derive from the lower ammonium to nitrate ratio. However, although genetic manipulation to diminish SA levels improved *pdx3* growth, it did not completely rescue growth and lesions became visible particularly when the *NahG* transgene was present. This is in contrast to full rescue upon ammonium fertilization or at elevated ambient temperature. To our surprise, *pdx3* lines were even more compromised when the SA receptor NPR1 was genetically removed, i.e. the *pdx3 npr1* double mutant. Therefore, the signaling response, even from a basal level of SA, is beneficial to *pdx3* lines that suffer a c/N imbalance, assisting growth at 22°C. Although we demonstrate that SA can alter the vitamin B_6_ profile, the changes (predominantly PM) do not overlap with those of loss of function *pdx3* mutants (increased PMP, decreased PLP). Thus, the increase in SA in *pdx3* lines may be a protective strategy that is consequential to the PMP:PLP imbalance leading to increased N assimilation that lowers the ammonium to nitrate content. Importantly at 28°C, we saw a shift in metabolism that revealed a higher proportion of C compounds even in wild type, which in *pdx3* may balance the c/N imbalance at 22°C. This observation would be in accordance with the recent report that PIF4-mediated thermomorphogenesis is dependent on sufficient sugar supply, reflected by a higher C status and signaled by trehalose-6-phosphate (T6P) inhibition of the KIN10 kinase subunit of SnRK1 (Hwang et al., 2019). The latter would otherwise (low sugar) phosphorylate PIF4 leading to its degradation (Hwang et al., 2019). We conclude that the thermomorphogenesis program is intact in *pdx3*.

## CONCLUSION

Overall, in the absence of PDX3, our data suggests that metabolic remodeling occurs as a function of the alteration of vitamin B_6_ homeostasis and plants are triggered into the N-gear scavenging endogenous ammonium that compromises NR levels, an effect that is exaggerated by a deficit in PLP-dependent serine biosynthesis. The defects are bypassed when plants are fed with exogenous ammonium or increased temperature (Figure 8) due to a different metabolic equilibrium under these conditions that is not reliant on PDX3. Thus, PDX3 could undergo negative selection in breeding crops that are overly dependent on ammonium-enriched fertilizers, such as during the green revolution. However, modern and future approaches that strive for the use of less (ammonium-based) fertilizers will likely reveal its indispensability and requirement for N management, particularly when nitrate is a source of N. Moreover, plants that depend on ammonium fertilization are likely compromised in the SA-mediated responses, as SA biosynthesis is reduced under this condition. Furthermore, PDX3 may be an important contributor in breeding for future climates with increased CO_2_ without having to increase ammonium fertilization. Our study suggests that PDX3 provides a surveillance mechanism for PMP:PLP ratios implying that these vitamers are sensed. It will be exciting in the future to unravel how this occurs.

## MATERIAL AND METHODS

### Plant material and growth conditions

*Arabidopsis thaliana* (Columbia ecotype), *pdx3-3* (SALK_054167C) and *pdx3-4* (GK-260E03), complementing lines *pdx3-3 35S::PDX3_1* and *pdx3-4 35S::PDX3_1* were described in (Colinas et al., 2016), independent complementing lines *pdx3-3 35S::PDX3_2* and *pdx3-4 35S::PDX3_2* were also included here. The autoimmune lines *bon1-1* described in (Hua et al., 2001), *snc1-1* described in (Li et al., 2001) were donated by Prof. Jian Hua (Cornell University). The plant line expressing the *NahG* transgene described in (Lawton et al., 1995) was donated by Dr. Christiane Nawrath and Prof. Philippe Reymond (University of Lausanne). The *sid2-1* line described in (Nawrath and Metraux, 1999) was donated by Prof. Roman Ulm (University of Geneva). The *npr1-2* line described in (Cao et al., 1997) was obtained from the European Arabidopsis Stock Center (N3801, NASC). For metabolic profiling, glutamate dehydrogenase activity assay, nitrate reductase activity assay, nitrate measurement, gene expression, and protein level analysis, plants were grown on soil (Einheitserde, Patzer classic ton Kokos) with a composition of 25% clay, 45% wheat peat, 15% brown peat, 15% coconut fiber pH 5.8. Before use, the soil was sterilized for 1 h at 60°C and supplemented once with a 2 ml l^-1^ solution of entomopathogenic nematodes (Andermatt, Traunem) and 2 tablespoons l^-1^ of *Bacillus thuringiensis israelensis* (Andermatt, Solbac). Plants were either watered with tap water (Geneva Switzerland; -) or supplemented every 9-10 days with either a 50 mM solution of potassium chloride (+KCl), potassium nitrate (+KNO_3_) or ammonium nitrate (+NH_4_NO_3_) in equal volumes and grown under a 16 h photoperiod (long-day) at 120-190 μmol photons m^-2^ s^-1^ generated by fluorescent lamps at 22°C and 8h darkness at 18°C or constant temperatures of 28°C with 60% relative humidity and ambient CO_2_ up to 21 days after germination. Rosette leaves were harvested 3 h after the onset of illumination for RT-qPCR, metabolite profiling and glutamate dehydrogenase activity assay, 3-6 h after the onset of illumination for nitrate content and HPLC, and 7 h after the onset of illumination for immunochemical analyses. For phenotypic comparison of plants grown under elevated CO_2_ (HC, 3000 ppm) or ambient CO_2_ (LC, 380 ppm), a 12-h photoperiod (120 μmol photons.m^-2^ s^-1^) at 22°C combined with 12 h of darkness at 18°C was used. For free ammonium measurement and serine supplementation, seeds were sown on plates containing modified MS medium containing no ammonium (Sigma-Aldrich M2909), 0.05% MES, 0.01% myo-inositol, and 0.55% agar. L-serine (100 μM) was added from a 0.1M filter sterilized stock solution.

### Generation of double mutants

The *pdx3-3 NahG*, and *pdx3-3 sid2-1* double mutants were generated using pollen from a *pdx3-3* plant to fertilize flowers of *NahG* and *sid2-1* plants. The *pdx3-4 NahG* and *pdx3-4 sid2-1* were generated in the same way by using the pollen of a *pdx3-4* plant. The *npr1-2 pdx3-3* double mutant, on the other hand, was generated using pollen from a *npr1-2* plant to fertilize flowers of a *pdx3-3* plant. The success of the cross was validated by the absence of a *pdx3*-like leaf phenotype of the F1 plants (due to the presence of one wild type copy of PDX3). The F2 plants were screened for homozygosity of the respective mutations by analyzing the phenotype frequency of their F3 offspring and by genotyping. The presence of the T-DNA insertion in *pdx3-3, pdx3-4*, or the *NahG* transgene was verified by PCR analysis of genomic DNA using oligonucleotides reported in Supplemental Table S1. The presence of a point mutation in *sid2-1* and *npr1-2* were verified by PCR analysis of genomic DNA using oligonucleotides reported in Supplemental Table S1, followed by Sanger sequencing of the PCR fragment (Microsynth AG).

### Immunochemical analyses

Five volumes of extraction buffer (50 mM Tris-HCl pH 8.5, containing 10 mM EDTA, 0.1% Triton X-100 (v/v), 10% glycerol and 1% (v/v) complete plant protease inhibitor cocktail (P9599, Sigma-Aldrich)) was added to frozen and ground plant material. After brief homogenization, samples were centrifuged for 15 min at 10,000 *g* at 4°C and the supernatant was collected. The protein concentration was determined by the Bradford method (Bradford, 1976). Samples were then separated by 10% SDS-PAGE loading 5 μg of total protein per lane. For western blot analysis, the proteins were transferred onto nitrocellulose membranes using the iBlot system (Invitrogen) for a total of 8 min applying 20 V for 1 min followed by 23 V for 4 min and finally 25 V for the remainder of the time. The membranes were stained with Ponceau S (0.1% Ponceau S in 5% acetic acid) to confirm protein transfer before removing the stain by washing in Tris buffered saline + 0.1% tween (TBS-T). The following primary antibodies and dilutions were used: α-PDX3 as described in (Colinas et al., 2014) 1:3000; α-NR (AS08310, Agrisera), 1:10000; α-ACTIN-2 as loading control (A0480, Sigma-Aldrich), 1:10000. The secondary antibody Goat Anti-Rabbit IgG (H + L)-HRP (1706515, Bio-rad) was used in combination with α-PDX3 at a dilution of 1:3000 and α-NR at a dilution of 1:10000. For α-ACTIN-2 the secondary antibody Goat Anti-Mouse IgG (H + L)-HRP conjugate (1706516, Bio-rad) was used at a dilution of 1:5000. The immunoblot analysis of PDX3 and ACTIN-2 was performed using a SNAP i.d. 2.0 system (Millipore) as described in (Colinas et al., 2014). For NR analysis, blots were blocked in 5% non-fat milk dissolved in TBS-T for 1 h at 22ºC with agitation followed by incubation in the primary antibody (in blocking reagent) for 1 h at 22ºC with agitation. The antibody solution was removed and the blot was washed 1×15 min and 3×5 min with TBS-T at 22ºC with agitation. Thereafter, blots were incubated in secondary antibody (in blocking reagent) for 1 h at 22ºC with agitation and washed as described above. Chemiluminescence was detected using ECL SuperBright (AS16 ECL-S, Agrisera) for NR and Western Bright ECL (K-12045, Advansta) for PDX3 and ACTIN-2 and captured using an Amersham Imager 680 system (GE Healthcare).

### Nitrate and free ammonium measurements

The method is based on (Zhao and Wang, 2017). Briefly, for extraction, 10 vol of deionized water was added to frozen and ground plant material, samples were homogenized and heated at 100°C for 30 min. The samples were centrifuged for 10 min at 15,000 *g* and the supernatant was decanted. A standard curve of 10-120 mg l^-1^ KNO_3_ was used. For the reaction, 10 μl sample or standard was mixed with 40 μl of 5% (w/v) salicylic acid (in H_2_SO_4_) and incubated at room temperature for 30 min. Thereafter, a yellow color complex was revealed after addition of 950 μl of 8% (w/v) of NaOH, 200 μl of the mixture was transferred to a 96-well plate and the OD_410_ was measured using a Synergy2 microplate reader (BioTek). Free ammonium in leaves was determined by the indophenol-blue reaction as described in (Scheiner, 1976) with adaptations. For extraction, 1 ml of 100 mM HCl was added to 100 mg frozen and ground plant material, samples were homogenized and 500 μl of chloroform was added. The samples were shaken for 15 min at 4°C and centrifuged for 10 min at 12,000 *g* at 8°C. The supernatant was decanted and transferred to a 1.5 ml Eppendorf tube containing 50 mg of acid-washed activated charcoal thoroughly vortexed and centrifuged for 5 min at 20,000 *g* at 8°C. The supernatant was decanted once more, transferred to a fresh tube and centrifuged again for 10 min at 20,000 *g* at 8°C and the supernatant decanted again. A standard curve of 0-100 μM (NH_4_)_2_SO_4_ was used. For the reaction, equal volumes of sample or standard and 100 mM HCl were mixed. In a 96-well plate, 20 μl of the sample mixture was added to 100 μl of solution I (1% (w/v) phenol and 0.005% (w/v) sodium nitroprusside) followed by the addition of 100 μl solution II (1% (w/v) sodium hypochlorite, 0.5% (w/v) sodium hydroxide). The plate was sealed with parafilm and incubated at 37°C for 30 min and OD_620_ was measured using a Synergy H1 microplate reader (BioTek).

### Gene expression analysis by RT-qPCR

RNA was extracted using the RNA NucleoSpin Plant kit (Macherey-Nagel) according to the manufacturers’ instructions. Reverse transcription (RT) was performed using 0.5-1 μg total RNA as template and Superscript II (Invitrogen) according to the instructions with the following modifications: stock oligo(dT)_15_ primers (C1101, Promega) concentration was 50 ng/μl and 0.5 μl Superscript II enzyme was used per reaction. qPCR was performed in 384-well plates on an Applied Biosystems QuantStudio 5 qPCR-System (Thermo Fisher Scientific) using PowerUp SYBR Green master mix (A25743, Applied Biosystems) and the following amplification program: 10 min denaturation at 95°C followed by 40 cycles of 95°C for 15 s and 60°C for 1 min. The data from nitrogen supplementation experiments was analyzed using the comparative cycle threshold method (2^−ΔCT^ or 2^−ΔΔCT^) normalized to the reference gene *ACT2* (At3g18780) and *UBC21* (At5g25760), whereas only *UBC21* was used in other experiments. Oligonucleotide pairs used are indicated in Supplemental Table S1. The two primer pairs for *GDH2* and *ATL31* gave similar results in independent experiments using the same dataset.

### Glutamate dehydrogenase activity

The animating (NADH-dependent) activity of glutamate dehydrogenase (GDH) was determined as described in (Turano et al., 1996) except that the extraction buffer consisted of 100 mM Tris-HCl pH 7.6, containing 1 mM MgCl_2,_ 1 mM EDTA and 14 mM β-mercaptoethanol.

### Vitamin B_6_ analysis by HPLC

Vitamin B_6_ profiling was performed as described in (Colinas et al., 2016) with the following changes: two separate extractions were performed with 15 vol and 8 vol of 50 mM ammonium acetate (pH 4), respectively, and a 50 μl injection volume was used for a single run per extract.

### Gas exchange measurements

Col-0 and *pdx3* mutants were grown under 12 h light (100–150 μmol photons m^2^ s^-1^) at 22°C and 12 h dark cycles at 18°C to promote vegetative growth at ambient air. For measuring the oxygen sensitivity of photosynthesis, oxygen concentrations at 2%, 20%, and 40% were generated as oxygen/nitrogen mixes by a gas-mixing system (Vögtlin instruments). The plants were acclimated for 20 min and the net photosynthetic rate was measured at 400 μl l^-1^ CO_2_ using an LI-6400XT (LiCor) photosynthesis analyzer.

### Metabolite profiling

For each analysis, six experimental replicates of Col-0, *pdx3-3* and *pdx3-4* were grown under long-day conditions. For the analysis of N supplementation, plants were either watered normally or supplemented every 9-10 days with a 50 mM solution of either KNO_3_ or NH_4_NO_3_ as indicated and harvested when they were 21 days old. The material of 4 to 6 plants was pooled and ground in liquid nitrogen using a mortar and a pestle. For data on the effect of high temperature, plants were subjected to a constant temperature of 28°C or to a temperature of 22°C during the photoperiod and 18°C during darkness and harvested when they were 12 days old and 14 days old, respectively. The material of 12 plants was pooled and ground in liquid nitrogen using a mortar and a pestle. The resulting powder was weighed and stored at - 80°C to be used for GC-MS as described in (Lisec et al., 2006) with peak annotation based on libraries of authentic standards (Kopka et al., 2005).

### SA treatment

Rosette leaves of 19-20 days old soil-grown Col-0 plants were sprayed equally with a solution of 0.005% Silwet L77 only as a mock treatment or containing either 2 mM SA (247588, Sigma-Aldrich). Treatment was started 0.5 h after the onset of light. Whole rosette leaves were harvested in liquid nitrogen in triplicate (pools of 5-8 plants) in a time-series of 3 h, 12 h, 24 h, and 48 h after treatment. All samples were harvested during the photoperiod.

### Nitrate reductase activity

For NR activity measurements with the purified protein the procedure was adapted from Sigma quality control tests for product N7265 based on (Gilliam et al., 1993). Arabidopsis NR2 (NIA2, At1g37130, ≥ 0.5 units of NR per vial) expressed and purified from *Pichia pastoris* was purchased from Sigma-Aldrich (N0163). The lyophilized protein was resuspended in 50 μl of 57 mM potassium phosphate at pH 7.5 and was used directly or stored at -80°C. Before the start of the assay the NR enzyme was diluted 100x in 57 mM potassium phosphate at pH 7.5 (≥ 0.01 units/μl). To circumvent the overlap in absorbance between PMP (82890, Sigma-Aldrich) and the NADH coenzyme, the latter was replaced with its analog, 3-acetylpyridine-adenine dinucleotide (APADH, MBS682175 MyBiosource). The reaction was carried out in a 96 well plate in 57 mM potassium phosphate pH 7.5 containing 0.005 mM FAD, 10 mM potassium nitrate, 0.1 mM APADH, 0 to 1 mM PMP and was started by the addition of 50 μl of NR (≥ 0.005 units added) for a total reaction volume of 200 μl. Alternatively, the reaction mixture without potassium nitrate was incubated for 25 min at 25°C before starting the reaction with the addition of 20 μl of 100 mM potassium nitrate (10 mM final concentration). The reaction was followed by measuring the absorbance at 363 nm at 25°C for 15 min using a plate reader (BioTek Synergy 2). For NR activity measurements on material extracted from Arabidopsis, the procedure described in (Colinas et al., 2016) was used with modifications. Whole rosette leaves of 21 days old plants grown under long day conditions were harvested in liquid nitrogen at 0 h (in the dark), 3 h, 6 h, and 12 h, after the onset of light and 2 h after the onset of the following period of darkness. For extraction, 50 μl of extraction buffer (250 mM Tris-HCl, pH 8.0 containing 1 mM EDTA, 5 μM FAD, 1 μM sodium molybdate, 3 mM DTT, 1% Triton-X-100, and 1% plant protease inhibitor cocktail (P9599 Sigma-Aldrich)) was added to 25 mg of frozen and ground plant tissue. After brief homogenization (samples kept on ice), samples were centrifuged 15 min at 13,000 *g* at 4°C and the supernatant was collected. When necessary, to further clear the extract from cell debris, the collected supernatant was centrifuged once more for 5 min at 13,000 *g* at 4°C. Using 96 well plates, 10 μl of plant extract or standard was added to 70 μl of 100 mM sodium phosphate pH 7.4 containing 40 mM sodium nitrate, and the reaction was started with the addition of 20 μl of 1 mM NADH (0.2 mM final concentration) for a total reaction volume of 100 μl. The reaction was incubated in the dark at 25°C, and was stopped immediately at 15-17 min, 30 min, and 45 min after the start of the reaction by adding 50 μl of stop solution (1:1 mixture of 1% (w/v) sulfanilamide and 0.05% (w/v) napthylethylenediamine). The absorbance at 540 nm was measured in a Synergy 2 microplate reader (BioTek).

### Analysis software

Data rendering and statistical analysis was performed using GraphPad Prism version 8.3.0 for Windows, GraphPad Software, San Diego, California USA, www.graphpad.com. Image rendering was performed using Inkscape 0.92.4 https://www.inkscape.org. Protein quantification was performed using ImageJ https://imagej.nih.gov/ij/. Photo color was modified for homogeneity using https://www.pixelmator.com/pro/.

## ACKNOWLEDGMENTS

This work was supported by the Swiss National Science Foundation (Grants 31003A-141117/1 and IZLIZ3_183193 to TBF) and the University of Geneva (TBF). MC was supported by a short term EMBO fellowship (ASTF 485-2014). APMW and ME acknowledge funding under Germany’s Excellence Strategy EXC-2048/1, Project ID 390686111. We thank Karolina Vogel for helping with the ammonia determination.

## AUTHOR CONTRIBUTIONS

PS, ME, MC, LR-S, TBF carried out experimental work, contributed tools and analyzed data. APMW and ARF analyzed data. TBF supervised the research & wrote the paper.

---

**Supplemental Figure S1.**
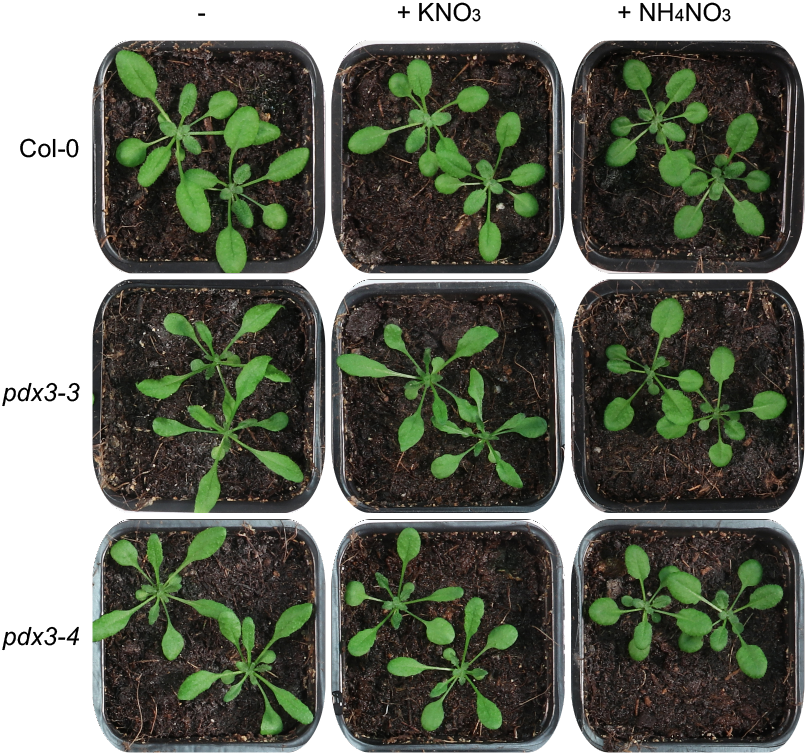
Phenotype of rosette leaves of *pdx3* compared to wild type. Photographs of wild type (Col-0) and *pdx3* lines grown on unfertilized (-) and either potassium nitrate (+ KNO_3_), ammonium nitrate (+ NH_4_NO_3_) fertilized soil. The plants are 21 days old and were grown on soil under a 16 h photoperiod (120-160 μmol photons m-2 s-1) at 22°C and 8 h darkness at 18°C and were watered either with water alone (-) or a 50 mM solution of the indicated compound every 9-10 days.

**Supplemental Figure S2.**
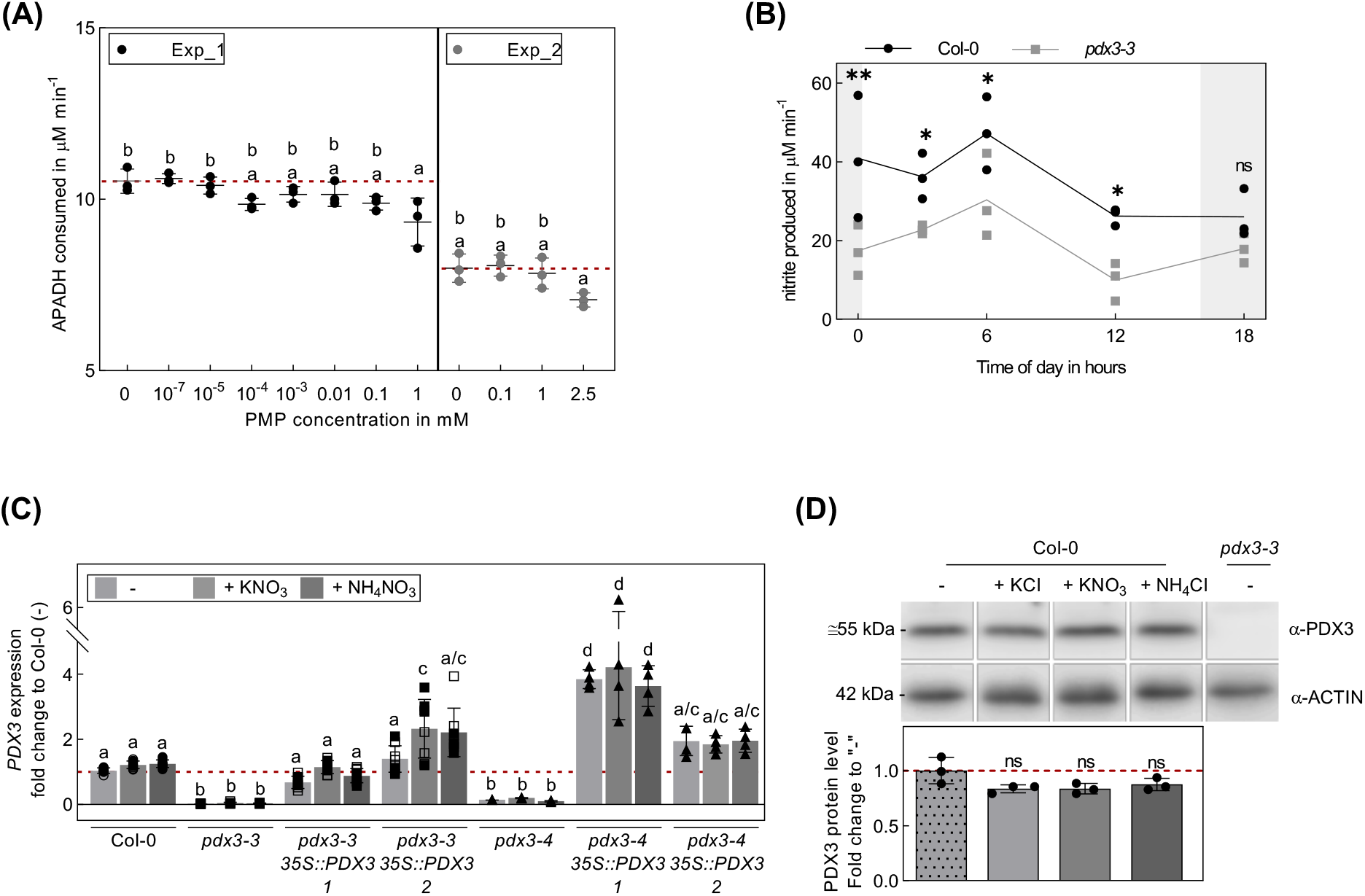
Nitrate reductase activity and *PDX3* expression as a function of PMP and N fertilization, respectively. (A) Activity of recombinant nitrate reductase in the presence of PMP, shown as rate of APADH (NADPH substitute) consumption in the presence of 0-2.5 mM PMP at pH 7.5 and 25°C. The data represents the mean ± SD of three technical and two experimental replicates (Exp_1 and Exp_2). Statistical analysis was performed using ordinary one-way ANOVA with Sidak’s multiple comparison test. (B) Nitrate reductase activity in rosette leaves of wild type and *pdx3-3* plants grown on unfertilized soil under a 16 h photoperiod (120-160 μmol photons m-2 s-1) at 22°C and 8 h darkness harvested at 0 h (in the dark before the onset of light), 3 h, 6 h, 12 h (all in the light), and 18 h (2 h after onset of darkness). The data represents the mean ± SD of 1 experimental and 3 biological replicates. Statistical analysis was performed using a two-tailed Student’s unpaired *t*-test with Col-0 as control (nsp>0.05, *p≤0.05, and **p≤0.005. (C) Relative expression of *PDX3* in *pdx3* and complementing lines compared to wild type (Col-0) grown on unfertilized (-) and either potassium nitrate (+ KNO_3_) or ammonium nitrate (+ NH_4_NO_3_) fertilized soil. Data represents the mean ± SD across 2 experimental replicates, (either open or filled symbols) except for *pdx3-4* and corresponding complementing lines, with 4 biological replicates each. Statistical analysis was performed using an ordinary one-way ANOVA with Tukey’s multiple comparisons test for the transcript (different letters indicate p≤0.05). (D) Protein levels of PDX3 in wild type (Col-0) and *pdx3-3* (as a control) grown on unfertilized soil and wild type grown on either potassium chloride (+ KCl), potassium nitrate (+ KNO_3_) or ammonium chloride (+ NH_4_Cl) fertilized soil. Data represents the mean ± SD of 3 biological replicates. Statistical analysis was performed using a two-tailed Student’s unpaired *t*-test using condition (-) as control (nsp>0.05). Plants were grown as in (B) and watered with water alone (-) or a 50 mM solution of the indicated compound every 9-10 days.

**Supplemental Figure S3.**
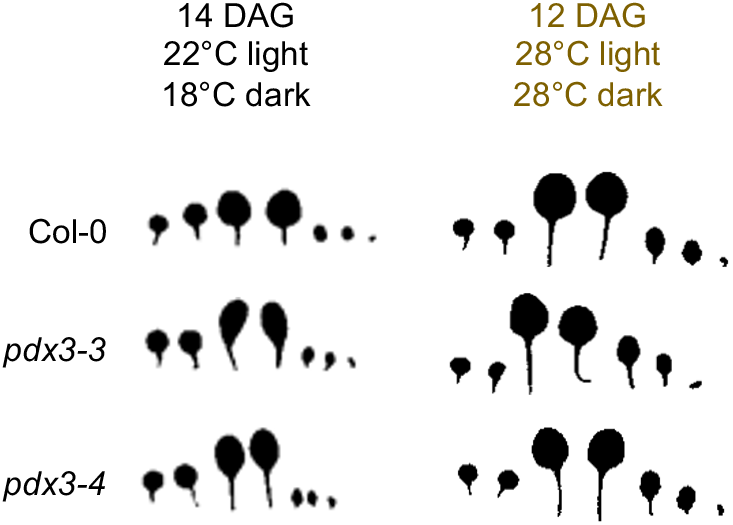
Photograph of the leaves of wild type (Col-0) and *pdx3* lines grown up to 14 days after germination (DAG) under the standard temperature of 22°C compared to 12 DAG under 28°C. In these conditions and developmental stage, the number of true leaves (five) is equal.

**Supplemental Table S1.**
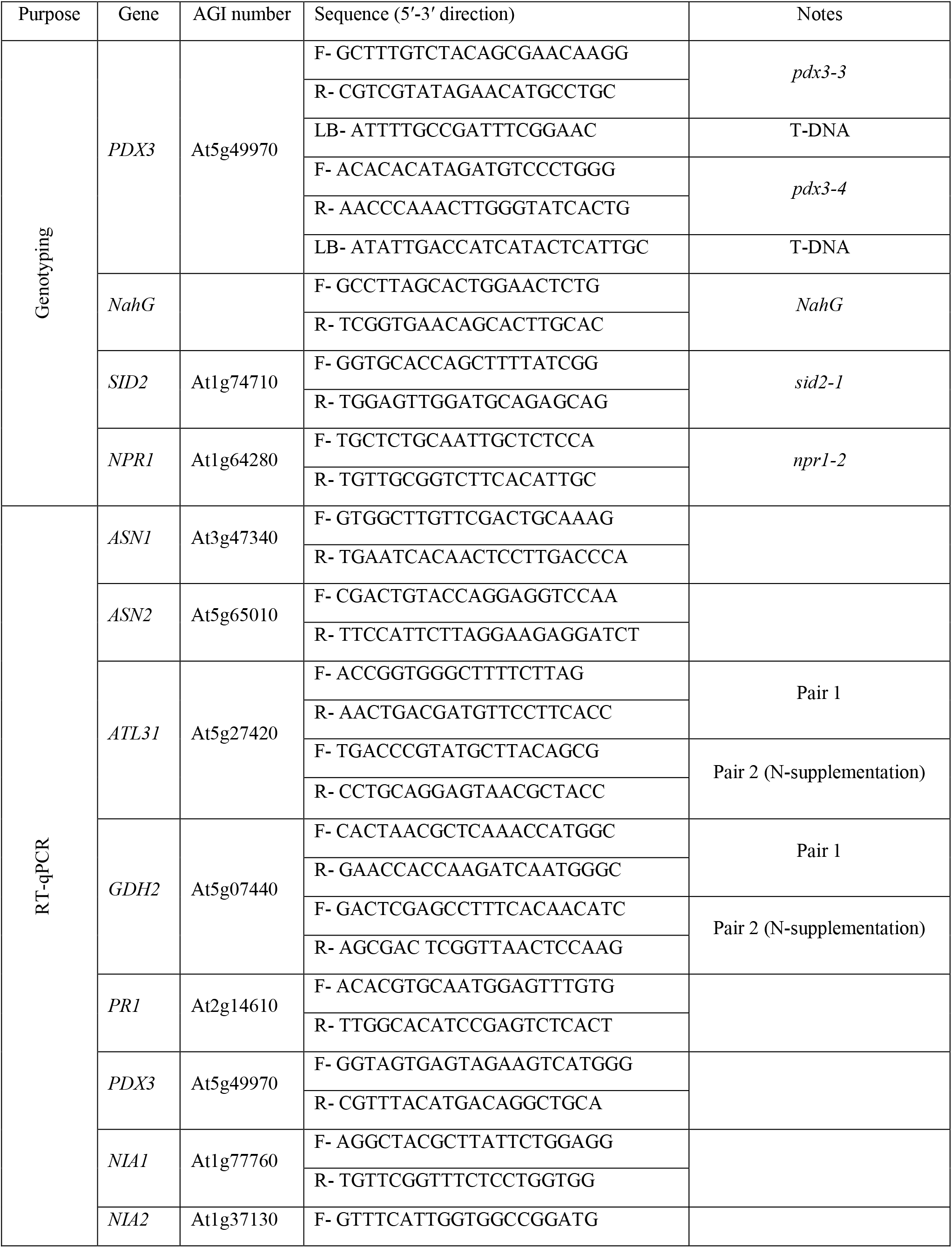
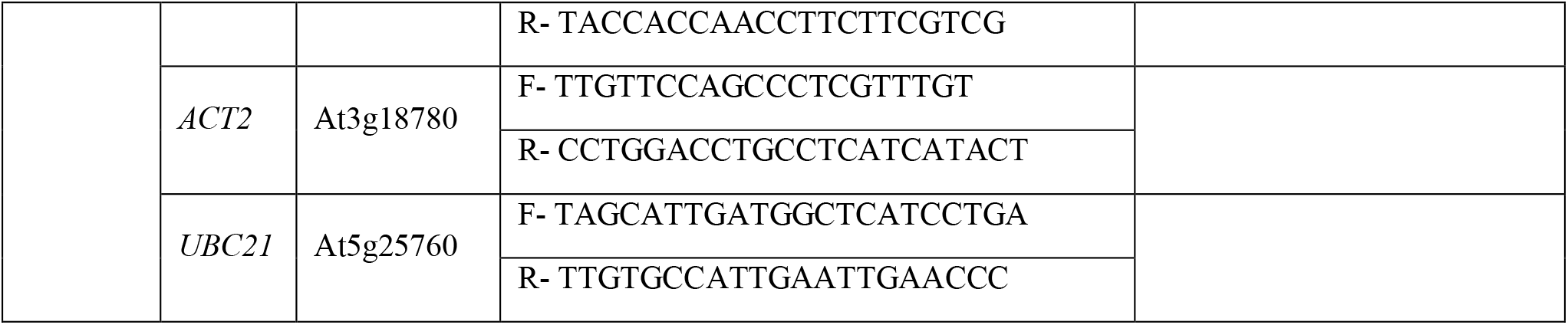
Oligonucleotides used in this study. Forward (F), reverse (R), left border (LB).

